# K^+^ uptake and root-to-shoot allocation in *Arabidopsis* require coordination of nitrate transporter1/peptide transporter family member NPF6.3/NRT1.1

**DOI:** 10.1101/674903

**Authors:** Xian Zhi Fang, Xing Xing Liu, Ya Xing Zhu, Jia Yuan Ye, Chong Wei Jin

**Author notes:** These authors contributed equally to this work (Xian Zhi Fang, Xing Xing Liu).

## Abstract

K^+^ and NO_3_^-^ are the major forms of potassium and nitrogen that are absorbed by the roots of most terrestrial plants. In this study, we observed that the close relationship between NO_3_^-^ and K^+^ homeostasis was mediated by nitrate transporter1 (NRT1.1) in *Arabidopsis*. The *nrt1.1* mutants lacking NRT1.1 function showed disturbed K^+^ uptake and root-to-shoot allocation, especially under K^+^-limited conditions, and had a yellow-shoot sensitive phenotype on K^+^-limited medium. The K^+^ uptake and root-to-shoot allocation of these mutants were partially rescued by expressing NRT1.1 in the root epidermis-cortex and central vasculature by using *Sultr1;2* and *PHO1* promoters, respectively. Furthermore, two-way analysis of variance based on the K^+^ content in *nrt1.1-1*/*akt1, nrt1.1-1*/*hak5-3, nrt1.1-1*/*kup7*, and *nrt1.1-1*/*skor-2* double mutants and their corresponding single mutants and wild-type plants revealed physiological interactions between NRT1.1 and K^+^ channels located in the root epidermis-cortex and central vasculature. Taken together, these data suggest that the expression of NRT1.1 in the root epidermis-cortex coordinates with K^+^ uptake channels to improve K^+^ uptake, whereas its expression in the root central vasculature coordinates with the channels loading K^+^ into the xylem to facilitate K^+^ allocation from the roots to the shoot.

## Introduction

Potassium (K^+^) is essential for plant growth and development, and it determines the yield and quality of crops in agriculture production (Wang & Wu, 2013). However, the concentration of soluble K^+^ in most soils is relatively low, which often limits plant growth (Maathuis, 2009). Crop production is increased by applying a large amount of potassic fertilizers to agricultural fields, but about one-half of the applied fertilizers is unavailable to plants and left behind as residues in soils, consequently leading to environmental contamination (Meena *et al*., 2016). Thus, improving the K^+^ utilization efficiency of plants by thoroughly understanding the molecular mechanisms of K^+^ transport and regulation is urgently required. Therefore, in the past few decades, researchers have been focusing on identifying K^+^ channels and transporters as well as their regulation mechanisms in plants.

In *Arabidopsis thaliana*, 71 K^+^ channels and transporters have been identified and categorized into three channel families (shaker, tandem-pore K^+^, and Kir-like) and three transporter families (KUP/HAK/KT, HKT, and CPA families) (Wang & Wu, 2010). Among them, the shaker inward K^+^ channel AKT1 and the KUP/HAK/KT K^+^ transporter HAK5 have been characterized as the two major components that mediate K^+^ uptake from the external environment into root cells, but they operate at different K^+^ levels (Pyo *et al*., 2010; Wang & Wu, 2013). AKT1 functions in plant K^+^ uptake over a wide range of K^+^ concentrations. Comparatively, HAK5 shows a high-affinity K^+^ transport activity (Gierth *et al*., 2005). After its uptake into root epidermal cells, K^+^ is then distributed to different organs or tissues of a plant. An *Arabidopsis* shaker-like outward-rectifying K^+^ channel, SKOR, was first identified to facilitate K^+^ secretion from the stelar cells to the xylem sap in the roots, which is a critical step for long-distance K^+^ transport from the roots to the shoots (Gaymard *et al*., 1998). Recently, *At*KUP7, one of the KT/HAK/KUP family members, was functionally characterized as a novel K^**+**^ transporter participating in both K^+^ uptake by roots from the growth medium and K^+^ release into the xylem for its allocation from roots to shoots, especially under K^+^-limited conditions (Han *et al*., 2016). However, the uptake affinity for K^+^ of KUP7 is considerably lower than that of *At*HAK5 (Wang & Wu, 2017).

In addition to the several aforementioned K^+^ channels and transporters, many other nutrients such as Na^+^, Ca^2+^, and N also remarkably affect K^+^ homeostasis in plants, particularly the association of N with K^+^, which has been explored for a long time because N is the mineral element that is required in the greatest quantity by most plants and is the most widely used fertilizer nutrient in crop production (Fageria & Baligar, 2005; Wang & Wu, 2013; Shin, 2017). Since the 1960s, physiological studies have revealed a close relationship between NO_3_^-^ and K^+^ with regard to uptake and translocation (Zioni *et al*., 1971; Blevins *et al*., 1978; Barneix & Breteler, 1985; Drechsler *et al*., 2015). However, how these two nutrients coordinate with each other in the transport pathways has not yet been clearly explored at the molecular level. Interestingly, lines of evidence have shown that NRT1.5 from the nitrate transporter1 (NRT1)/peptide transporter (PTR) transporter family (NPF) functions as a component in controlling K^+^ allocation in plants (Lin *et al*., 2008; Drechsler *et al*., 2015; Li *et al*., 2017; Du *et al*., 2019), which was initially identified as a pH-dependent bidirectional NO_3_^-^ transporter and is mainly expressed in the root pericycle cells mediating NO_3_^-^ release into the xylem vessels (Lin *et al*., 2008). In this study, we showed that the loss of another nitrate transporter1 member NRT1.1 in *nrt1.1* mutants also led to the development of a more serious K-deficiency phenotype under low-K stress. Further physiological and genetic evidences revealed that both the uptake and root-to-shoot allocation of K^+^ in plants require NRT1.1 in plants. However, NRT1.1 acts as a coordinator rather than a K^+^ channel/transporter in the above two K^+^ nutrition issues, which could depend on its transport activity for NO_3_^-^. Our findings suggest that the modification of NRT1.1 activity in crops by using biological engineering approach might be a promising way for simultaneously improving the utilization efficiencies of K and N fertilizers in agriculture production.

## Material and Methods

### Plant material

The mutants *chl1-5* (Huang et al., 1996), *nrt1.1-1* (salk_097431), *nrt1.2* (cs859605), *nrt2.1* (cs859604), *nrt2.2* (salk_043543), *nrt2.5* (GK-213H10.06), *akt1* (salk_071803), *hak5* (salk_130604), *kup7* (cs805085), and *skor* (cs2103489) and the transgenic plants *pNRT1.1::NRT1.1-GFP* and *pNRT1.1::NRT1.1-GUS* (cs6513) are in a Columbia (Col-0) background, whereas the *chl1-6* (cs6154) and *nrt2.4* (cs27332) lines are in a Landsberg *erecta* (*Ler*) background. The seeds of the lines *chl1-5* and *pNRT1.1::NRT1.1-GFP* were kindly gifted by Dr. Philippe Nacry (Biochimie et Physiologie Moléculaire des Plantes, Montpellier, France). The double mutants *nrt1.1-1/akt1, nrt1.1-1/hak5-3, nrt1.1-1/kup7*, and *nrt1.1-1/skor-2* were generated by crossing *nrt1.1-1* with *akt1, hak5-3, kup7*, and *skor-2*, respectively. The mutants were verified using the primers listed in Table S1.

### Plant culture

The seeds were surface-sterilized and sown on basal agar medium containing 1% sucrose (w/v) and 0.9% (w/v) agar. The complete nutrients in the basal agar medium had the following composition: 1.125 mM Ca(NO_3_)_2_, 500 μM CaCl_2_, 500 μM MgSO_4_, 750 μM NaH_2_PO_4_, 375 μM (NH_4_)_2_SO_4_, 25 μM Fe-EDTA, 10 μM H_3_BO_3_, 0.5 μM MnSO_4_, 0.5 μM ZnSO_4_, 0.1 μM CuSO_4_, and 0.1 μM (NH_4_)_6_Mo_7_O_24_, pH 6.5. After 4 d of germination, the seedlings were used in studies, and the final concentrations of K^+^ and NO_3_^-^ were adjusted using KCl and Ca(NO_3_)_2_, as described in the figure legends.

### Phenotypic analyses

After 8 d of growth in agar medium containing 2 mM K^+^ or 0.05 mM K^+^, the phenotype and fresh weight of plants were recorded. The chlorophyll of fresh leaves was extracted with 80% (v/v) acetone and was colorimetrically measured according to Choi *et al*. (2014). In addition, the chlorophyll fluorescence was analyzed using a pulse amplitude-modulating imaging fluorometer (IMAGING-PAM; Germany). Briefly, after a 20 min dark-adaption, the leaves were arranged on the fluorometer, and the maximum quantum efficiency (Fv/Fm) and electron transport rate through PSII (ETR) were monitored.

### Measurement of K concentrations

The harvested plants were separated into shoots and roots. Samples were dried at 75°C for 48 h, and then wet-digested as described by Luo *et al*. (2012). The digests were diluted with ultrapure water, and the K concentrations were analyzed using MP-AES (Agilent Technologies; USA).

### Green fluorescent protein and β-glucuronidase expression analysis

The NRT1.1-green fluorescent protein (GFP) expression in the roots of *pNRT1.1::NRT1.1-GFP* plants was observed using an epifluorescence microscope (excitation, 488 nm; emission, 525–550 nm; Ni-U; Nikon). Histochemical assay of β-glucuronidase (GUS) expression in the roots of *pNRT1.1::NRT1.1-GUS* plants was performed as described by Fang *et al*. (2016), and the distribution and intensity of the blue product were observed under a microscope.

### Complementation of *nrt1*.*1-1* with *pSultr1;2::NRT1*.*1* and *pPHO1::NRT1*.*1*

The 2 kb genomic fragments upstream of the *PHO1* and *Sultr1;2* initiation codons were, respectively, PCR-amplified on genomic DNA from Arabidopsis by using the primers listed in Table S1. Subsequently, the above two amplified products were cloned upstream of the NRT1.1 coding region by using the pCAMBIA1300 vector. The constructs were transformed in *nrt1.1-1* plants by using the Agrobacterium-mediated floral dip method. Homozygous T4 transgenic plants were used.

### Measurement of NO_3_^-^ and K^+^ and Rb^+^ uptake

Net fluxes of NO_3_^-^ and K^+^ were measured in the roots by using a non-invasive microelectrode ion flux measurement system (ipa-2; AE, USA). After 3 d of preculture in the nutrient medium containing 2 mM or 0.05 mM K^+^, the flux of NO_3_^-^ and K^+^ in each roots section was monitored using NO_3_^-^-selective and K^+^-selective microelectrodes, as described by Hawkins *et al*. (2008) and Li *et al*. (2012), respectively. For Rb^+^ uptake, the 4-d-old seedlings were grown on complete nutrient medium containing 2 mM or 0.05 mM K^+^ replaced with Rb^+^ for 12 h, and then these plants were harvested and used for Rb concentration analysis.

### Transcription analysis

Total RNA in roots was extracted using RNAisoPlus (TaKaRa, Otsu, Japan) and was treated with DNase I to remove genomic DNA contamination. The first-strand cDNA was synthesized from genomic DNA-removed RNA by using PrimeScript™ RT reagent kit (TaKaRa). Next, the transcript levels of the corresponding genes were measured using quantitative real-time PCR by using TB Green™ Premix Ex Taq™ II (TaKaRa). The primers used are listed in Table S1.

### RNA sequencing and functional enrichment analysis

Total RNA was extracted from the roots of Col-0 and *nrt1.1-1* plants by using the MagZol Reagent (Magen), according to manufacturer’s protocol. Sequencing libraries were prepared using NEBNext Ultra RNA Library Prep Kit for Illumina (NEB, USA), and the quality was assessed using the Agilent 2100 Bioanalyzer. Sequencing was performed on a Hiseq Xten (Illumina) at RIBOBIO (Guangzhou, China). Raw sequencing reads were quality controlled and trimmed using Trimmomatic tools and FastQC, and then the clean reads were aligned to the TAIR10 reference genome by using HISAT2. Significantly differentially expressed genes were assessed using an adjusted *P*-value threshold of <0.05 and |log_2_(fold change)| of >1 by using DEGseq. All the genes with significant differential expression were grouped based on k-means clustering and selected for subsequent GO analyses by using R package clusterProfiler.

### Grafting of Arabidopsis plants

Grafts between *nrt1.1-1* mutants and Col-0 plants were performed following the protocol described by Andersen *et al*. (2014). Briefly, seedlings to be grafted were grown on complete nutrient agar medium for 5 days. After cotyledons were removed, the seedlings were separated into scion and rootstock by using a sterilized blade. The scion was then grafted to the corresponding rootstock on new agar plates by using forceps under a stereomicroscope. The grafted seedlings were left undisturbed for another 5–7 days to form graft union. The adventitious roots emerging at or above the graft union were eliminated, and then the successful unified seedlings were used in the study as described in the legend of figure 4.

**Figure 1.**
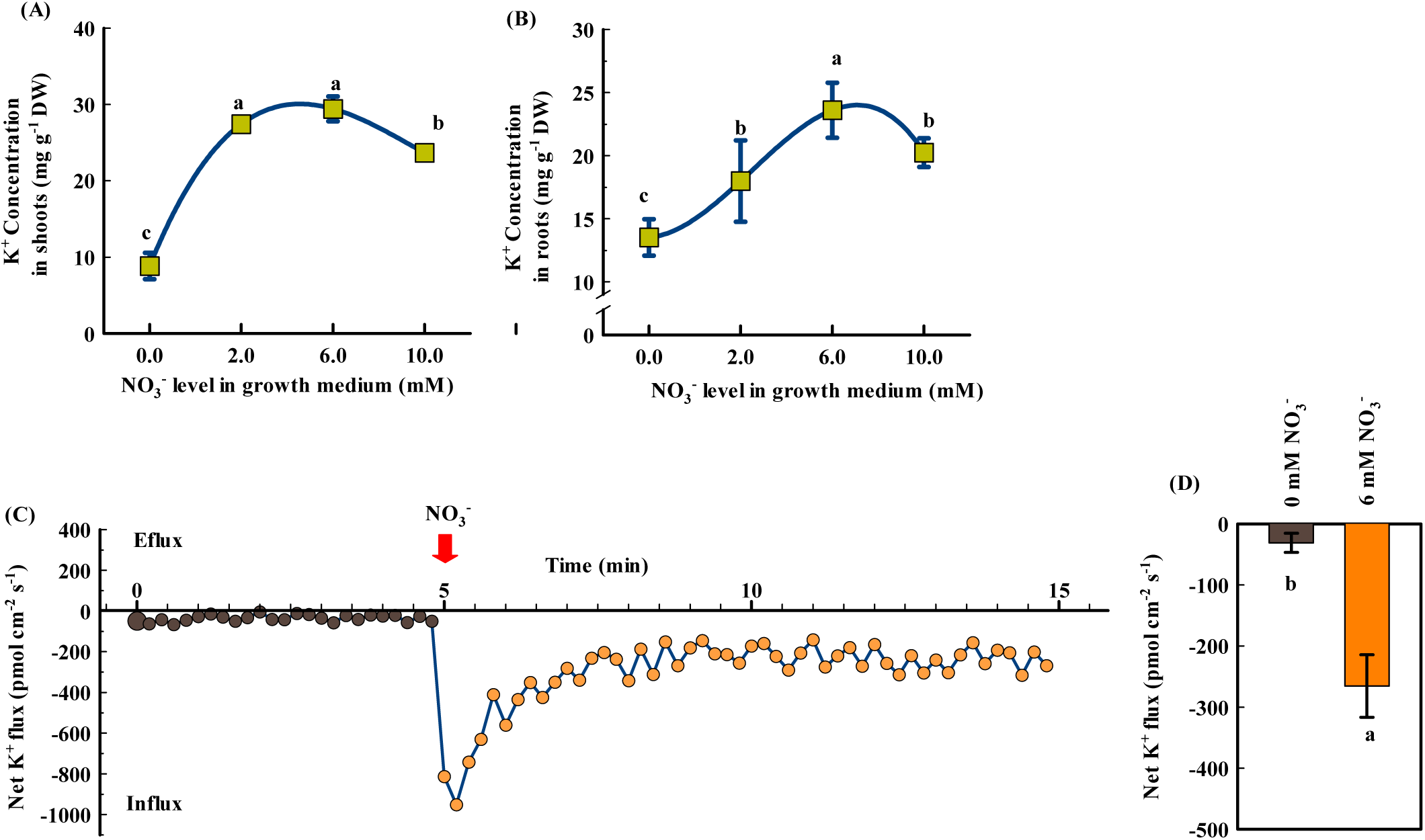
Increase of NO_3_^-^ supply improves K nutrition in *Arabidopsis* Col-0 plants. (A) and (B) The K concentration in the shoots and roots. After 4 d of germination on the basal agar media, as described in the Materials and Methods, the seedlings were transplanted to the basal agar media containing different NO_3_^-^ levels as indicated. The K concentrations were analyzed 8 d after seedling transfer. Bars represent the SD (n = 5 biological replicates). Different letters above bars indicate significant differences at *P* < 0.05 (LSD test). DW, dry weight. (C) Kinetics of net K^+^ flux in Col-0 roots in response to NO_3_^-^ addition. The net K^+^ flux in Col-0 roots was first measured for 5 min in the absence of NO_3_^-^, and then it was measured in the presence of 6 mM NO_3_^-^ (indicated by the red arrow). (D) The average values of root K^+^ fluxes calculated from Figure c. Bars represent the mean ± SD. Different letters above bars indicate significant differences at *P* < 0.05 (LSD test). DW, dry weight.

**Figure 2.**
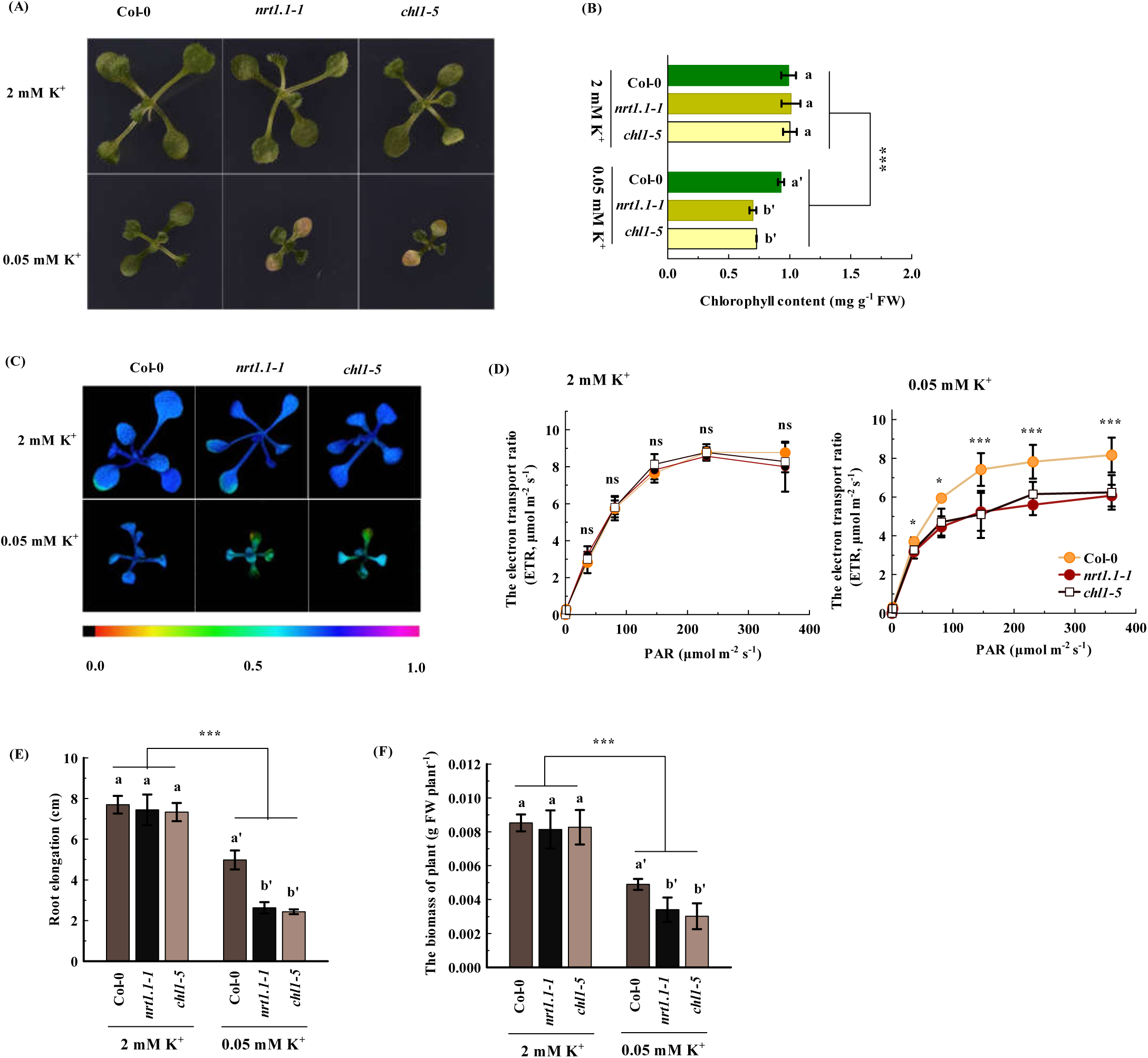
Loss of NRT1.1 function in *Arabidopsis nrt1.1* mutant leads to hypersensitivity to low-K stress. (A) Phenotypic photographs of plants; (B) chlorophyll content of leaves; (C) maximal quantum efficiency of PS; (D) electron transport rate (ETR); (E) root elongation; (F) biomass of plants. The 4-d-old seedlings of Col-0, *nrt1.1-1*, and *chl1-5* were transferred to an agar medium containing 2 mM or 0.05 mM K^+^, as described in the Materials and Methods section. The analyses were performed 8 d after seedling transfer. Bars represent the mean ± SD (n = 4–5). (B), (E), and (F) Different letters above bars indicate significant differences at *P* < 0.05 (LSD test); asterisks indicate a significant genotype by treatment interaction (*** *P* < 0.001, two-way ANOVA). (D) Asterisks show significant differences compared with the Col-0; ns, non-significant (* *P* < 0.05, *** *P* < 0.001, two-tailed Student’s *t*-test).

**Figure 3.**
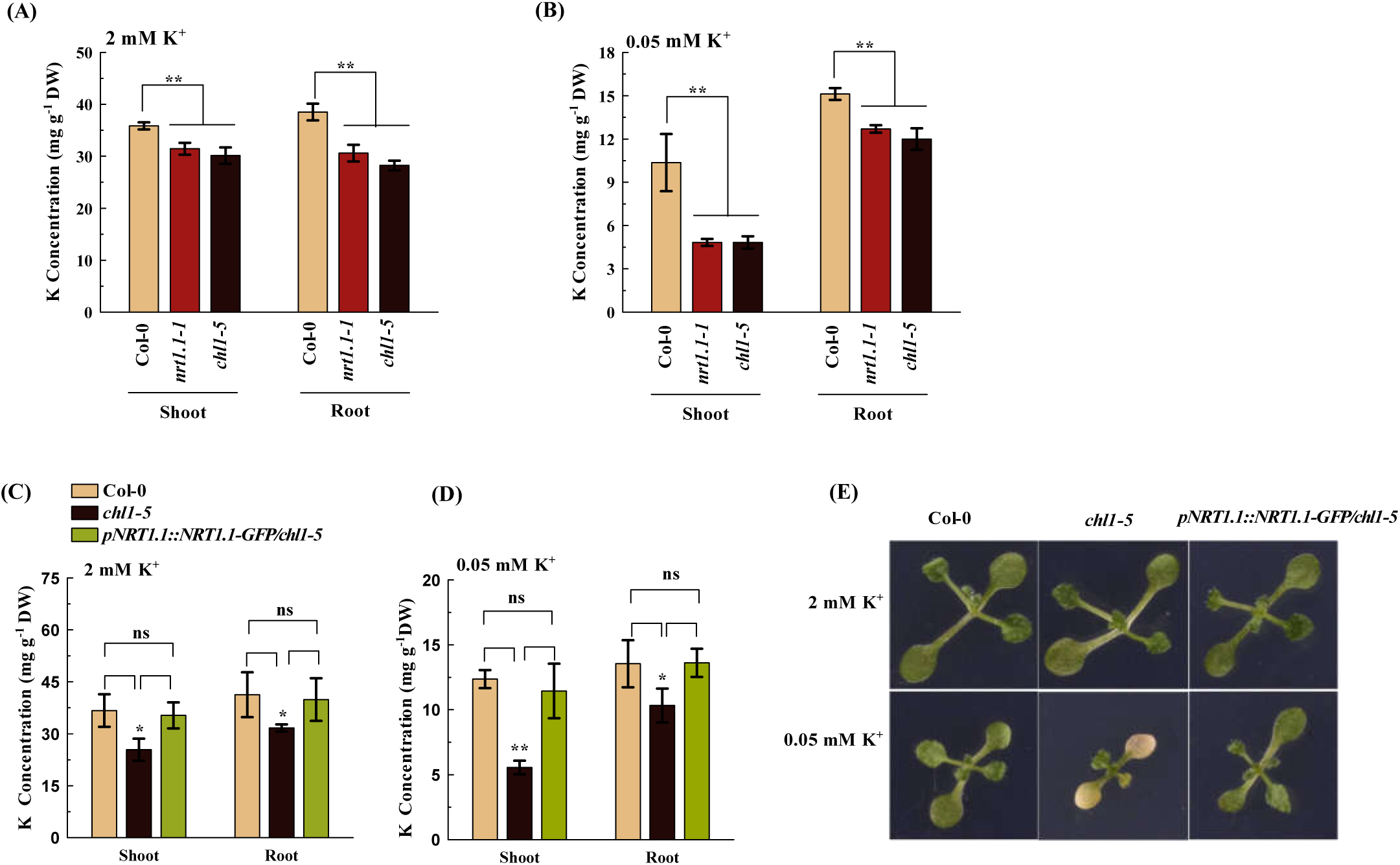
NRT1.1 is required for ensuring K nutrition in *Arabidopsis*. (A) and (B) Shoot and root K concentrations of Col-0, *nrt1.1-1*, and *chl-5* plants. (C) and (D) Shoot and root K concentrations of Col-0, *chl1-5*, and *pNRT1.1::NRT1.1-GFP* plants. (E) Phenotypic photographs of Col-0, *chl-5*, and *pNRT1.1::NRT1.1-GFP* plants. The 4-d-old seedlings were transferred to an agar medium containing 2 mM K^+^ or 0.05 mM K^+^, as described in the Materials and Methods section. The analyses were performed 8 d after seedling transfer. Bars represent the mean ± SD (n = 5). Asterisks indicate significant differences between genotypes; ns, non-significant (* *P* < 0.05, ** *P* < 0.01, two-tailed Student’s *t*-test). DW, dry weight.

**Figure 4.**
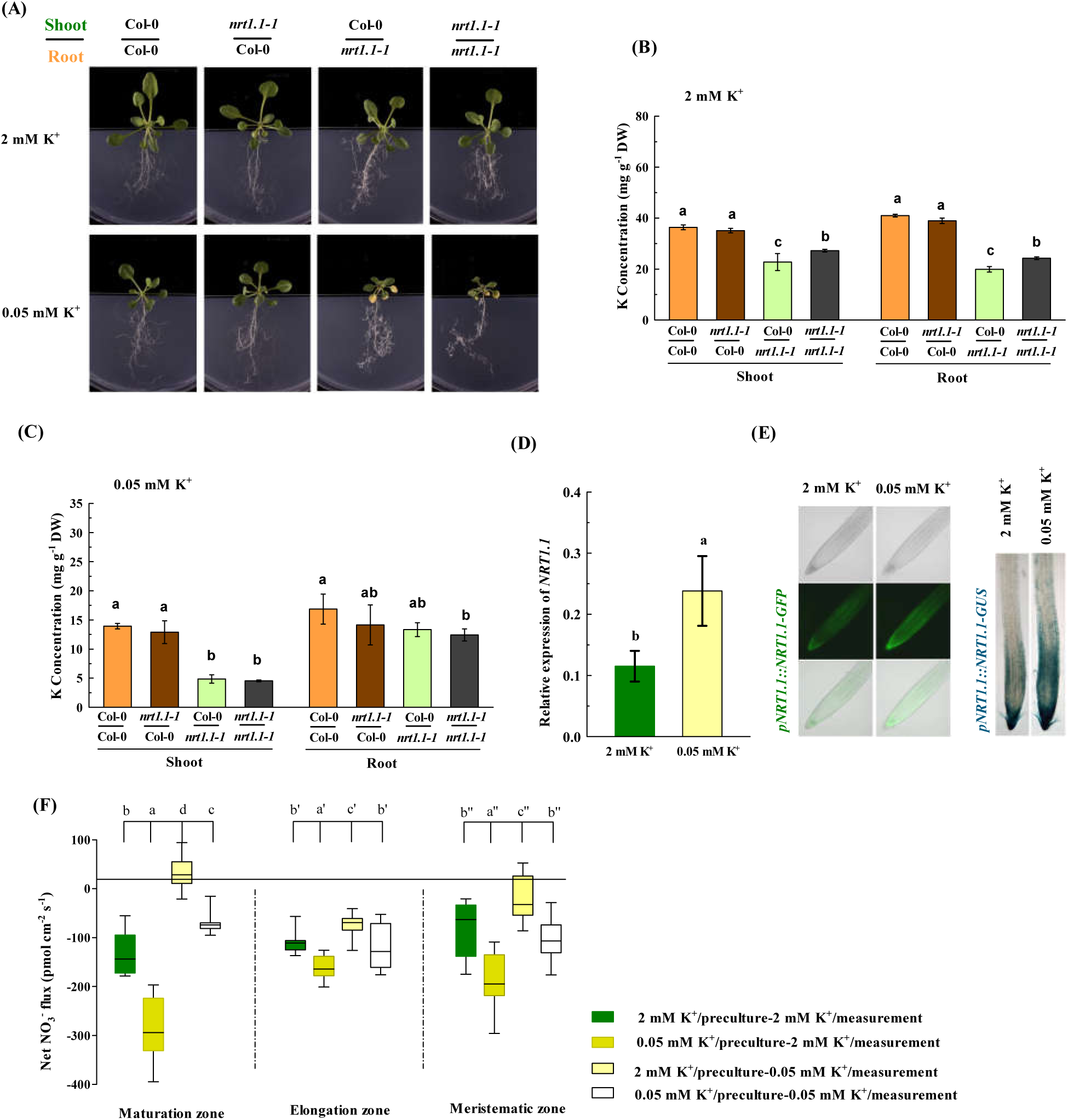
Improvement of K nutrition by NRT1.1 in *Arabidopsis* is a root behavior, and low-K stress stimulates NRT1.1 activity in the roots. (A) The phenotypic photographs; (B) and (C) the K concentrations of grafted plants. The unified seedlings were transferred to an agar medium containing 2 mM K^+^ or 0.05 mM K^+^. The analyses were performed 8 d after seedling transfer. (D) Relative expression of *NRT1.1*. (E) NRT1.1-GFP expression and GUS staining in *pNRT1.1::NRT1.1-GFP* and *pNRT1.1::NRT1.1-* GUS transgenic plants, respectively. (F) Average values of net NO_3_^-^ fluxes in the meristematic, elongation, and maturation zones of Col-0 roots. The 4-d-old seedlings were transferred to an agar medium containing 2 mM K^+^ or 0.05 mM K^+^, as described in the Materials and Methods section. The analyses were performed 5 d after seedling transfer. The relative expression levels were normalized to the levels of *UBQ10*. The negative value in (F) indicates net influx. Bars represent the mean ± SD (n = 5); different letters above bars indicate significant differences at *P* < 0.05 (LSD test). DW, dry weight.

### Yeast complementation assay

The protein coding sequence of NRT1.1 was constructed into the pDR196 vector by using primers listed in Table S1. Next, the pDR196-NRT1.1 vector was transformed into the yeast strain R5421 (*trk1Δ, trk2Δ*) by using the lithium acetate method (Ito *et al*., 1983). Yeast strain R757 was used as a positive control. The picked single yeast cells were precultured overnight at 30°C in 2 mL liquid YPDA medium containing 100 mM KCl. The cells were collected the following day and washed with ultrapure water three times. Subsequently, the yeast cells were resuspended in ultrapure water to an OD_600_ of 1, and 10-fold serial dilutions from OD_600_ of 1 to 10^−3^ were incubated on AP agar plates supplemented with different K^+^ concentrations for the experiments.

## Results

### NO_3_^-^ facilitates K^+^ uptake by roots

We first determined how the NO_3_^-^ supply affects the K levels in Arabidopsis wild-type (Col-0) plants. The presence of NO_3_^-^ in the growth medium increased the K levels in both roots and shoots compared with those in the NO_3_^-^-free treatment (Fig. 1a,b). However, a further increase of NO_3_^-^ supply over 6 mM decreased the K levels in plants. These results indicate that an appropriate NO_3_^-^ supply might favor the uptake of K^+^ by roots. Therefore, we calculated the amount of K^+^ absorbed per weight of roots and found that these values also increased along with the supply of NO_3_^-^ when the level was less than 6 mM (Fig. S1). To further evaluate how NO_3_^-^ influences K^+^ uptake by Col-0 roots, we used the non-invasive micro-test technology (NMT system) to measure the transmembrane K^+^ influx and efflux by root cells in a non-invasive manner (Li *et al*., 2012). Although the net K^+^ influx by root cells was observed in the NO_3_^-^-free medium (Fig. 1c), its average value was only around −30 pmol cm^-2^ s^-1^ (Fig.1d, a negative value indicates influx). Interestingly, addition of 6 mM NO_3_^-^ to the rooting media resulted in a sudden increase in the net K^+^ influx rate, which reached around −900 pmol cm^-2^ s^-1^ (Fig. 1c). Subsequently, the rate of transmembrane K^+^ influx gradually decreased to a relatively stable level, but the average value was still about 7-fold higher than that of the NO_3_^-^-free treatment (Fig. 1c,d). Taken together, these results suggest that the uptake of K^+^ by roots might require a coordination of NO_3_^-^ uptake.

### Resistance to K deficiency requires NRT1.1

As the uptake of K^+^ requires a coordination of NO_3_^-^ uptake in roots, we intended to determine whether the sensitivity of a plant to low K stress is also associated with the root NO_3_^-^ uptake. In *Arabidopsis*, six nitrate transporters, including NRT1.1, NRT1.2, NRT2.1, NRT2.2, NRT2.4, and NRT2.5, have been characterized to be involved in the root NO_3_^-^ uptake (Wang *et al*., 2012; Léran *et al*., 2014; Lezhneva *et al*., 2014). Here we found that, in the growth medium containing 6 mM NO_3_^-^, the two NRT1.1 knockout mutants *nrt1.1-1* and *chl1-5* exhibited severer leaf senescence than in the wild-type plants (Col-0) after 8 d of low K treatment (0.05 mM K), but these three plant lines did not show distinguishable phenotypes under sufficient K treatment (2 mM K; Fig. 2a) Considering that the decrease of chlorophyll content would affect the photosynthesis of plants, the maximum quantum efficiency of photosystem II (Fv/Fm) and photosynthetic electron transport rate (ETR) were also evaluated as indicators of the photosynthetic reactions in leaves. In the leaves of *nrt1.1-1* and *chl1-5* plants, the Fv/Fm was almost completely abolished (Fig. 2c), and the ETR also declined apparently compared with that in Col-0 plants under low-K conditions (Fig. 2d). Therefore, the *nrt1.1* knockout mutants had both shorter roots and less biomass than the Col-0 plants after low-K treatment (Figs 2e,f, S2). In addition, we examined a third NRT1.1-null mutant, *chl1-6*, the fresh weight and root elongation of which were also significantly reduced compared with those of its corresponding wild-type *Ler* under low-K conditions (Fig. S3a,b). Since NRT1.1 can act as a high-affinity as well as low-affinity system (Wang *et al*., 2012; Léran *et al*., 2014), the growth of these NRT1.1-null mutants was also evaluated in low-K growth medium containing 0.2 mM NO_3_^-^. However, the phenotypes of these mutants were indistinguishable from those of wild-type plants (Fig. S4a,b). These results suggest that NRT1.1 plays an important role in plants to confer resistance to K deficiency, but it requires a sufficient supply of NO_3_^-^ in the growth medium.

We next determined the roles of the other five NRTs in plant growth responses to low-K stress. Since NRT2.1, NRT2.2, NRT2.4, and NRT2.5 act as high-affinity NO_3_^-^ transporters, the growth of these NRTs-null mutants was evaluated in a growth medium containing 0.2 mM NO_3_^-^. However, the fresh weight and root elongation of *nrt2.1, nrt2.2, nrt2.4*, and *nrt2.5* mutants were similar to those of the corresponding wild-type plants in both low-K and sufficient-K treatments (Fig. S5). Although NRT1.2 is a low-affinity NO_3_^-^ transporter (Huang *et al*., 1999), the *nrt1.2* mutant showed similar fresh weight and root elongation as those of wild-type plants in both low-K and sufficient-K growth medium containing 6 mM NO_3_^-^ (Fig. S5e,j). These results indicate that NRT1.2, NRT2.1, NRT2.2, NRT2.4, and NRT2.5 do not act like NRT1.1 when conferring resistance to K deficiency.

### NRT1.1 is required for homeostasis of K in plants

We next emphasized on how NRT1.1 affects K nutrition in plants. In both sufficient-K and low-K treatments, the K levels were significantly lower in *nrt1.1-1, chl1-5*, and *chl1-6* mutants than in their crossponding wild-type plants (Figs 3a,b, S3c,d), indicating that the lack of NRT1.1 function disturbed the K homeostasis in plants. To provide further support for the role of NRT1.1 in K homeostasis, we measured the K levels in the transgenic plants *pNRT1.1::NRT1.1-GFP*, which are in the *chl1-5* mutant background (Krouk *et al*., 2010). In both sufficient-K and low-K treatments, the *pNRT1.1::NRT1.1-GFP* transgenic plants contained higher K levels than those in the *chl1-5* mutants and similar K levels as those in Col-0 plants (Fig. 3c,d). Therefore, phenotype analysis showed that complementation of NRT1.1 conferred the transgenic plants tolerance to K^+^ deficiency, as evidenced by the similar chlorophyll content, root elongation, and fresh weight as those in Col-0 plants (Figs 3e, S6).

As described above, the uptake of K^+^ by roots requires a coordination of NO_3_^-^ uptake. However, when the plants were grown in the medium containing low NO_3_^-^ (0.2 mM), no significant difference in both growth and K levels was noted between *NRT1.1-null* mutants and wild-type plants in both sufficient-K and low-K treatments (Fig. S4). The result suggested that the role of NRT1.1 in maintaining K homeostasis in plants depends on sufficient NO_3_^-^ supply. We also investigated the role of the other NRTs in maintaining K homeostasis. Nevertheless, the K levels in the roots and shoots of *nrt1.2, nrt2.1, nrt2.2, nrt2.4*, and *nrt2.5* mutants were not obviously different from those of their corresponding wild-type plants in sufficient-K or low-K treatments (Fig. S8). Thus, NRT1.1 might improve the K nutrition in a relatively specific manner.

### Root is the action point for NRT1.1 to improve K nutrition in plants, and low-K stress stimulates the activity of NRT1.1 in roots

NRT1.1 is expressed in both roots and shoots (Guo *et al*., 2001). Therefore, homografted (Col-0 scion/Col-0 stock and *nrt1.1-1* scion/*nrt1.1-1* stock) and heterografted (Col-0 scion/*nrt1.1-1* stock and *nrt1.1-1* scion/Col-0 stock) plants were generated to clarify whether either or both the roots and shoots are the action point for NRT1.1 to improve K nutrition in plants. The leaves of Col-0/*nrt1.1-1* and *nrt1.1-1*/*nrt1.1-1* plants developed severely chlorotic phenotype after 8 d of being grown on low-K medium (Fig. 4a). However, *nrt1.1-1*/Col-0 and Col-0/Col-0 did not exhibit any chlorotic symptom. This result suggested that the absence of the root-part function of NRT1.1 was primarily responsible for the sensitive symptom. This conclusion was further supported by the observation that K levels were clearly reduced in the shoots of the plants grafted with *nrt1.1-1* root stock in low-K treatment (Fig. 4c). Furthermore, we found that the K levels in both the shoots and roots of Col-0/*nrt1.1-1* and *nrt1.1-1*/*nrt1.1-1* plants were significantly lower than those in the plants grafted with Col-0 rootstocks in the sufficient K treatment (Fig. 4b). These results suggested that only the root part is the action point for NRT1.1 to improve K nutrition in plants.

The role of root NRT1.1 in improving plant K nutrition prompted us to investigate how NRT1.1 in roots responds to low-K stress. Real-time quantitative PCR analysis showed that low-K treatment clearly increased the expression of *NRT1.1* in roots (Fig. 4d), whereas the expression of the other five NO3^-^ uptake transporter genes was either decreased (*NRT2.1*) (Fig. S9a) or not significantly affected (*NRT1.2, NRT2.2, NRT2.4*, and *NRT2.5*; Fig. S9b-e), suggesting that low-K stress might have a specific effect on the action of NRT1.1. Consistent with gene expression, the NRT1.1-GFP protein expression indicated by GFP-associated fluorescence in the roots of *pNRT1.1::NRT1.1-GFP* transgenic plants was elevated in the low-K treatment than in the sufficient-K treatment (Fig. 4e). The result was confirmed by GUS staining of the roots of *pNRT1.1::NRT1.1-GUS* transgenic plants (Fig. 4e). These findings suggested that the activity of NRT1.1 could be up-regulated in response to low K^+^ stress. Thus, we analyzed the rate of net NO_3_^-^ fluxes at the surface of the maturation, elongation, and meristematic zones of Col-0 roots by using the NMT system. In all the three measured zones, the plants precultured with low-K medium had a higher rate of net NO_3_^-^ influx than in those precultured with sufficient-K medium in the same testing medium (Fig. 4f), providing a direct evidence that low-K stress stimulates NO_3_^-^ uptake activity, which should be associated with an up-regulation of NRT1.1 due to low-K stress.

### Both uptake and root-to-shoot allocation of K require a coordination of NRT1.1

We next determined how NRT1.1 improves K nutrition in plants by investigating its role in the root K uptake. The alkaline ion Rb^+^, the closest analog of K^+^, was first used to evaluate the K^+^ uptake by roots. The rates of root Rb^+^ uptake in *chl1-5* and *nrt1.1-1* mutants were considerably less than those in Col-0 plants in the medium containing either 2 mM Rb^+^ or 0.05 mM Rb^+^ (Fig. 5a), indicating that the loss of NRT1.1 function might disturb the uptake of K^+^ by roots. To further evaluate the contribution of NRT1.1 in root K^+^ uptake, we measured the root K^+^ fluxes of the above three plant lines. We found that, at the elongation and maturation zones, the rates of net K^+^ influx of *nrt1.1-1* and *chl1-5* mutants were less than 50% of those obtained in the Col-0 plants in both 2 mM K^+^ and 0.05 mM K^+^ testing medium. However, the net K^+^ influx rate in the meristematic zone did not differ significantly among these plants (Fig. 5b,c). These results suggest that the elongation and maturation zones, but not the meristematic zone, are the target regions for NRT1.1 to contribute to K^+^ uptake by root cells.

**Figure 5.**
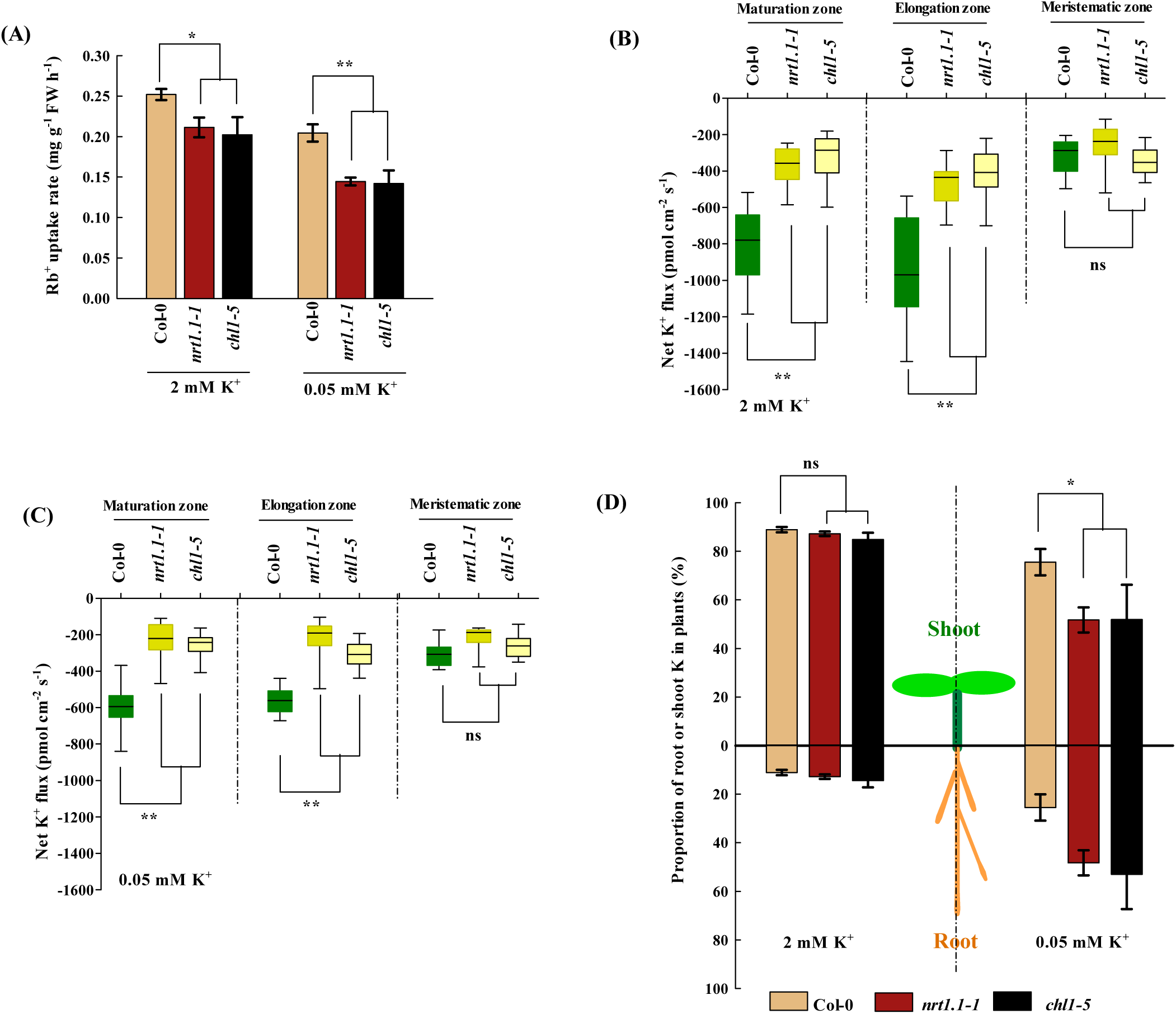
Both the uptake and root-to-shoot allocation of K^+^ require an action of NRT1.1 in *Arabidopsis*. (A) Rb^+^ uptake rate of roots. The 4-d-old seedlings were transferred to an agar medium containing 2 mM Rb^+^ or 0.05 mM Rb^+^ for 12 h, as described in the Materials and Methods section. Bars represent the mean ± SD (n = 5). (B) and (C) The net K^+^ fluxes in root. The 4-day-old seedlings were used to measure the net K^+^ fluxes in the meristematic, elongation, and maturation zones under 2 mM K^+^ or 0.05 mM K^+^ treatment. Negative indicates net influx; bars represent the mean ± SD (n = 15). (D) Proportions of K distributed in the shoots and roots. The 4-d-old seedlings were transferred to an agar medium containing 2 mM K^+^ or 0.05 mM K^+^, as described in the Materials and Methods section. The analyses were performed 8 d after seedling transfer. Bars represent the mean ± SD (n = 5); asterisks indicate significant differences compared with Col-0 plants; ns, non-significant (* *P* < 0.05, ** *P* < 0.01, two-tailed Student’s *t*-test).

Consistent with its role in NO_3_^-^ uptake, NRT1.1 was expressed in the epidermis and cortex of roots (Huang *et al*., 1996). However, NRT1.1 was also expressed in the root central vasculature (Remans *et al*., 2006), revealing its another function: allocating root NO_3_^-^ to shoot parts through xylem NO_3_^-^ loading in the root stele (Léran *et al*., 2013). This additional function suggests that NRT1.1 might also play a role in root-to-shoot allocation of K. Therefore, the rates of K distributed in the root and shoot were calculated. However, the proportion of K distributed in either shoots or roots did not differ between the Col-0 plants and *nrt1.1* knockout mutants when they were grown in sufficient-K medium (Fig. 5d). Interestingly, both *chl1-5* and *nrt1.1-1* mutants had less proportion of K distributed in the shoots, but greater proportion of K distributed in the roots compared with those in Col-0 plants grown in the low-K medium (Fig. 5d). These results indicated that a K-insufficient status of plants allows NRT1.1 to play the distinct role in root-to-shoot allocation of K.

### Action of NRT1.1 in favoring uptake and root-to-shoot allocation of K, respectively, depends on its specific expression in the epidermis-cortex and central vasculature of roots

As NRT1.1 is expressed in both the epidermis-cortex and central vasculature of roots (Huang *et al*., 1996; Remans *et al*., 2006), we intended to determine whether such expression pattern correlates with its dual roles in coordinating the uptake and root-to-shoot allocation of K^+^. To address this question, we generated transgenic plants expressing NRT1.1 under the control of the promoters of *Sultr1;2* and *PHO1* genes, respectively, in the *nrt1.1-1* mutant background. *Sultr1;2* is expressed in the root epidermis and cortex (Yoshimoto, 2002), whereas *PHO1* is expressed in the root central vasculature (Hamburger *et al*., 2002; Wege & Poirier, 2014). Therefore, the uses of these two promoters to drive *NRT1.1* gene in transgenic plants could, respectively, ensure NRT1.1 expression in the epidermis-cortex and central vasculature of roots, although their expression levels might not well match with those in the wild-type plants. The determination of K levels in two independent *pSultr1;2::NRT1.1* transgenic lines in comparison to those in Col-0 and *nrt1.1-1* plants revealed partial recovery of K to wild-type levels in the plants in both sufficient-K and low-K treatments (Fig. 6a,b). However, the calculation of the ratio of K distributed in the root and shoot showed that *pSultr1;2::NRT1.1* complementation in *nrt1.1-1* plants did not improve the root-to-shoot allocation of K (Fig. 6c). Therefore, the expression of NRT1.1 in the epidermis and cortex might only be responsible for improving the K uptake.

**Figure 6.**
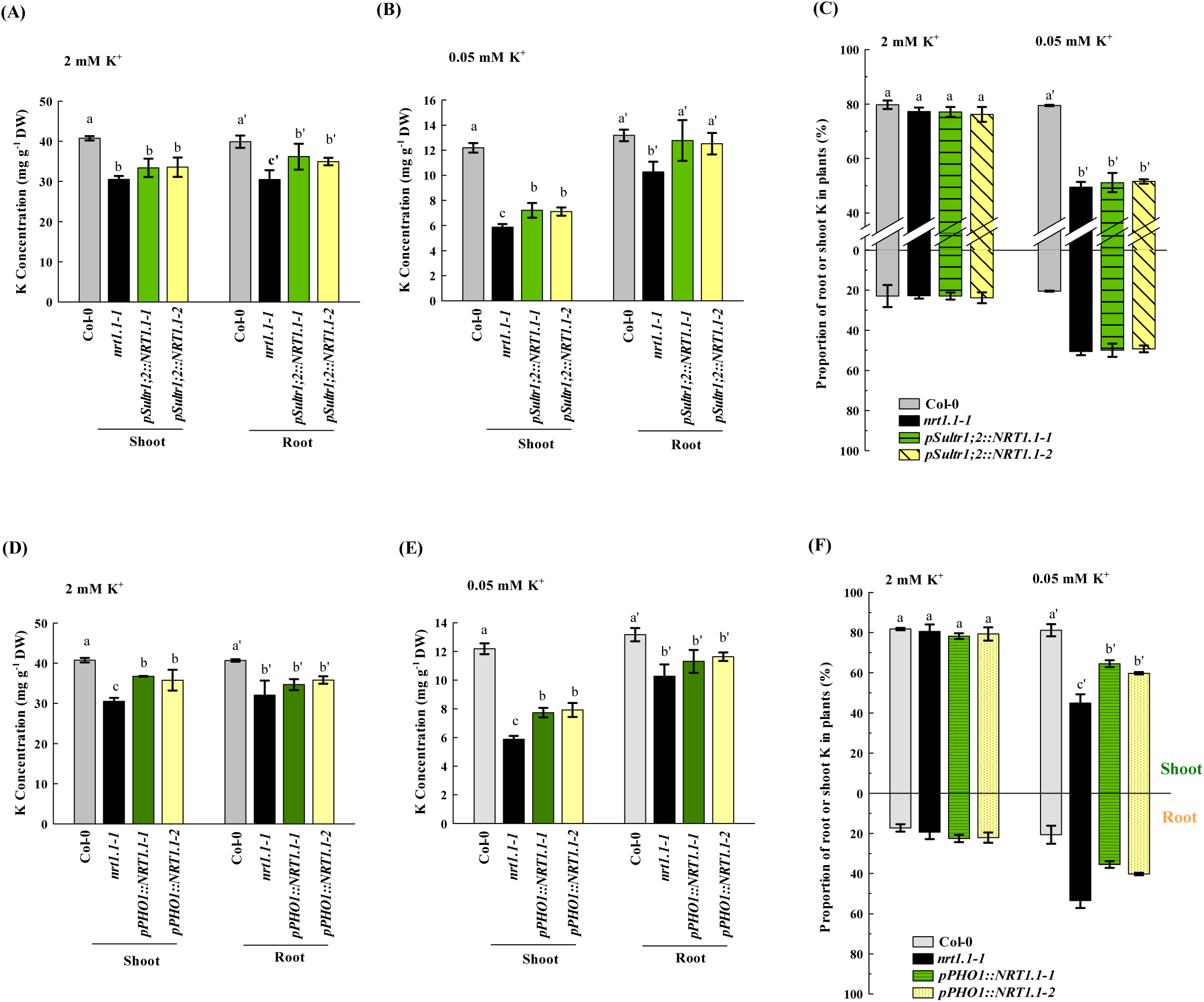
*NRT1.1* expression driven by *Sultr1;2* and *PHO1* promoters partially rescued the uptake and root-to-shoot allocation of K in *nrt1.1* mutants, receptively. (A) and (B) K concentrations and (C) distributions in Col-0, *nrt1.1-1*, and two independent *pSultr1;2:NRT1.1*-transformed *nrt1.1-1* lines (*pSultr1;2:NRT1.1-1/-2*). (D) and (E) K concentrations and (F) distributions in Col-0, *nrt1.1-1*, and two independent *pPHO1:NRT1.1*-transformed *nrt1.1-1* lines (*pPHO1:NRT1.1-1/-2*). The 4-d-old seedlings were transferred to an agar media containing 2 mM K^+^ or 0.05 mM K^+^ for 8 d, as described in the Materials and Methods section. Bars represent the mean ± SD (n = 5); different letters above bars indicate significant differences at *P* < 0.05 (LSD test).

The measurement of K levels in other two independent *pPHO1::NRT1.1* transgenic lines also showed partial recovery of K to Col-0 wild-type levels in the plants in both K treatments, with the recovery being more distinct in the shoots (Fig. 6d,e). Furthermore, the ratio of K distributed in the root and shoot revealed that the complementation of *pPHO1::NRT1.1* in *nrt1.1-1* plants increased the K transport from root to shoot (Fig. 6f), suggesting that the expression of NRT1.1 in the central vasculature of roots is responsible for supporting the root-to-shoot allocation of K.

### Action of NRT1.1 in favoring the uptake and root-to-shoot allocation of K, respectively, depends on the K^+^ channels in the epidermis-cortex and central vasculature of roots

Next, we investigated how the expression of NRT1.1 in the epidermis-cortex and central vasculature, respectively, coordinated the uptake and root-to-shoot allocation of K. In addition to a role in mediating nitrate transport across the plasmalemma, NRT1.1 was found to act as a singling component in regulating the expression of numerous genes in plants (Ho *et al*., 2009; Bouguyon *et al*., 2015; Wang *et al*., 2018). Therefore, the difference in whole-genome gene expression profiles between Col-0 plants and *nrt1.1* mutants in response to different K treatments were compared using RNA-seq. Gene ontology (GO) analysis revealed that the differentially expressed genes (DEGs) were involved in many physiological processes such as response to nutrient levels, response to reactive oxygen species, and response to salicylic acid (Fig. S10). However, among the genes associated with K uptake and trafficking, only the expression of *SKOR*, which mediates K^+^ loading into the xylem (Gaymard *et al*., 1998; Drechsler *et al*., 2015), was inhibited by NRT1.1 mutation in both sufficient-K and low-K treatments, whereas that of the other genes was either increased or not affected (Fig. 7a). This result cannot explain the decreased root K uptake due to the loss of function of NRT1.1. Therefore, we also investigated whether NRT1.1 acts a K^+^ channel. To functionally test the role of NRT1.1 in K^+^ transport activity in a heterologous expression system, we used the yeast (*Saccharomyces cerevisiae*) mutant R5421, lacking the main K^+^ uptake systems (*trk1Δ, trk2Δ*; Gaber *et al*., 1988; Han *et al*., 2016). However, NRT1.1 could not complement the growth retardation of R5421 under K deficiency (Fig. 7b), refuting the K^+^ transport activity of NRT1.1.

**Figure 7.**
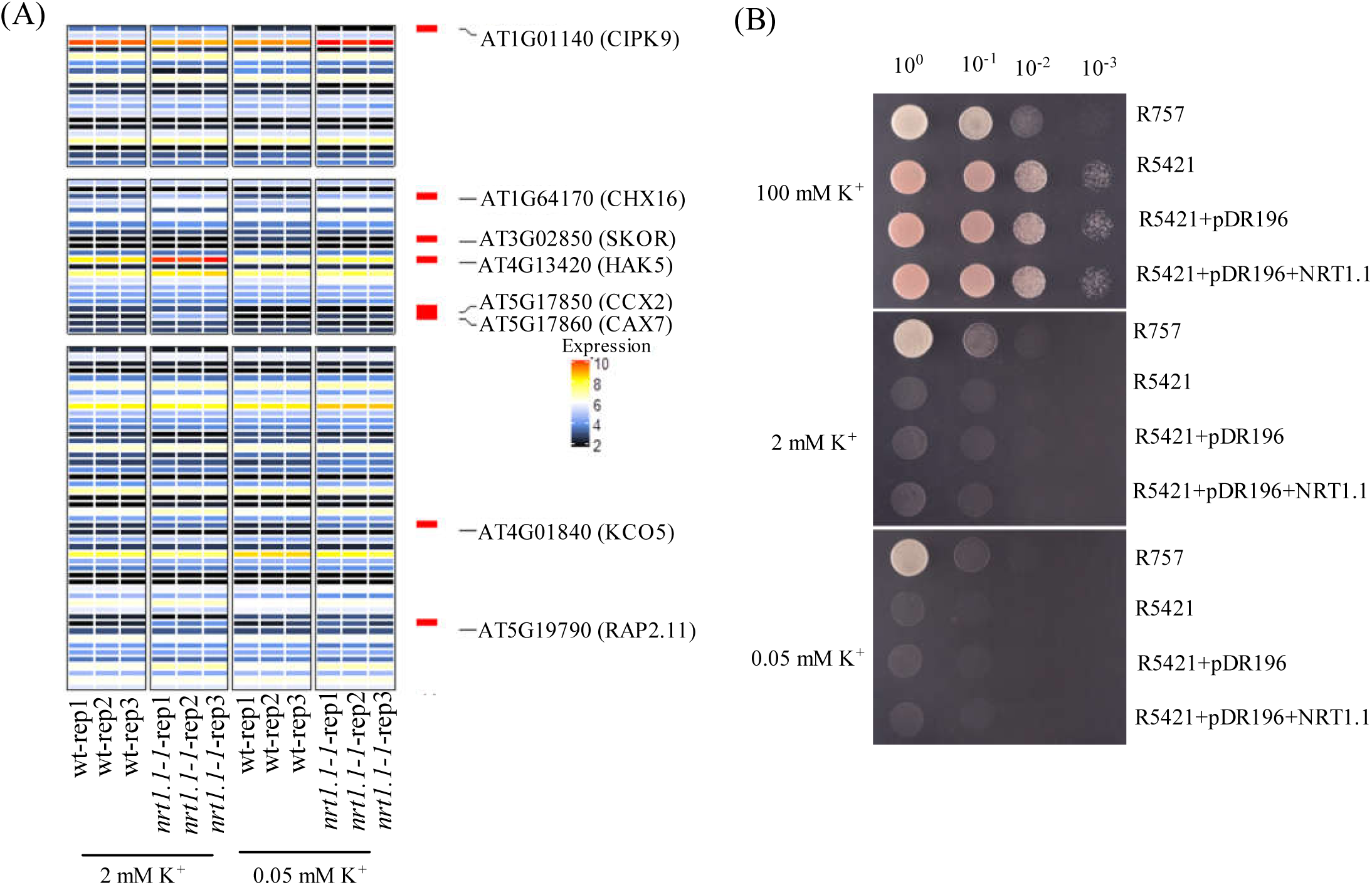
Both the regulation of gene expression and K^+^ transport activity might not explain the effect of NRT1.1 in improving K nutrition in *Arabidopsis*. (A) Heatmap analysis of genes enriched in the GO terms: K^+^ homeostasis, K^+^ transport, and other processes of K^+^ metabolism in the roots of *Arabidopsis* Col-0 and *nrt1.1-1* plants. The 4-d-old seedlings were transferred to a media containing 2 mM or 0.05 mM K^+^ for 5 d, and roots were collected to perform transcriptome analysis, as described in the Materials and Methods section. Red color represents significantly up- or down-regulated genes compared with those in the control group. Changes in response to K treatment with a *p*-value of <0.05 were selected for GO analysis. (B) Functional analysis of NRT1.1 in *Saccharomyces cerevisiae*. K^+^ uptake activity was analyzed in yeast R757 strain and the K^+^ import mutant strain R5421 (*trk1Δ, trk2Δ*), which were transformed with the expression constructs, as described in the Materials and Methods section. pDR196, empty vector.

Theoretically, the flux of anions across the membrane should be accompanied by an orthokinetic flux of cations so as to maintain the ionic balance, and this process might depend on the cooperation between anion transporters/channels and cation transporters/channels. In this context, the action of NRT1.1-mediated NO_3_^-^ flux across the plasmalemma might collaborate wih K^+^ channels. To test this hypothesis, we generated *nrt1.1-1*/*akt1, nrt1.1-1*/*hak5-3, nrt1.1-1*/*skor-2*, and *nrt1.1-1*/*kup7* double mutants by crossing the NRT1.1-null mutant *nrt1.1-1* with *akt1, hak5-3, skor-2*, and *kup7* mutants and measured K levels in these plants. AKT1, HAK5, and KUP7 are the three characterized components that contribute to K^+^ uptake in *Arabidopsis* roots, and the latter two channels only have a significant contribution in K uptake under K-limited condition (Lagarde *et al*., 1996; Gierth *et al*., 2005; Han *et al*., 2016). In addition, KUP7 was found to be involved in K^+^ loading into the xylem in addition to the SKOR channel (Han *et al*., 2016). Two-way analysis of variance (ANOVA) showed that the difference of K levels in the shoots and roots between *akt1* and *akt1/nrt1.1-1* mutants was significantly less than that between Col-0 and *nrt1.1-1* in both K treatments (Fig. 8a,b), suggesting that the reduced K levels due to the loss of NRT1.1 was associated with the action of AKT1. Similarly, two-way ANOVA based on the data obtained in Col-0, *nrt1.1-1, hak5-3, nrt1.1-1/hak5-3, kup7*, and *nrt1.1-1*/*kup7* plants showed that the reduced K levels due to the loss of NRT1.1 were also associated with HAK5 and KUP7 in low-K medium (Fig. 8c-e). These results indicated that NRT1.1 should interplay with the K uptake channels in the root epidermis-cortex to improve the plant K nutrition. Next, we determined whether the KUP7 and SKOR channels are required for the action of NRT1.1 in favoring root-to-shoot allocation of K under low K condition. Two-way ANOVA showed that the difference in the proportion of K distributed from the roots to shoots either between *skor-2* and *nrt1.1-1/skor-2* or between *kup7* and *nrt1.1-1*/*kup7* was significantly less than that between Col-0 and *nrt1.1-1* (Fig. 8f,h). These results indicate that the action of NRT1.1 in favoring root-to-shoot allocation of K depends on the K^+^ channels that mediate K^+^ loading into the xylem.

**Figure 8.**
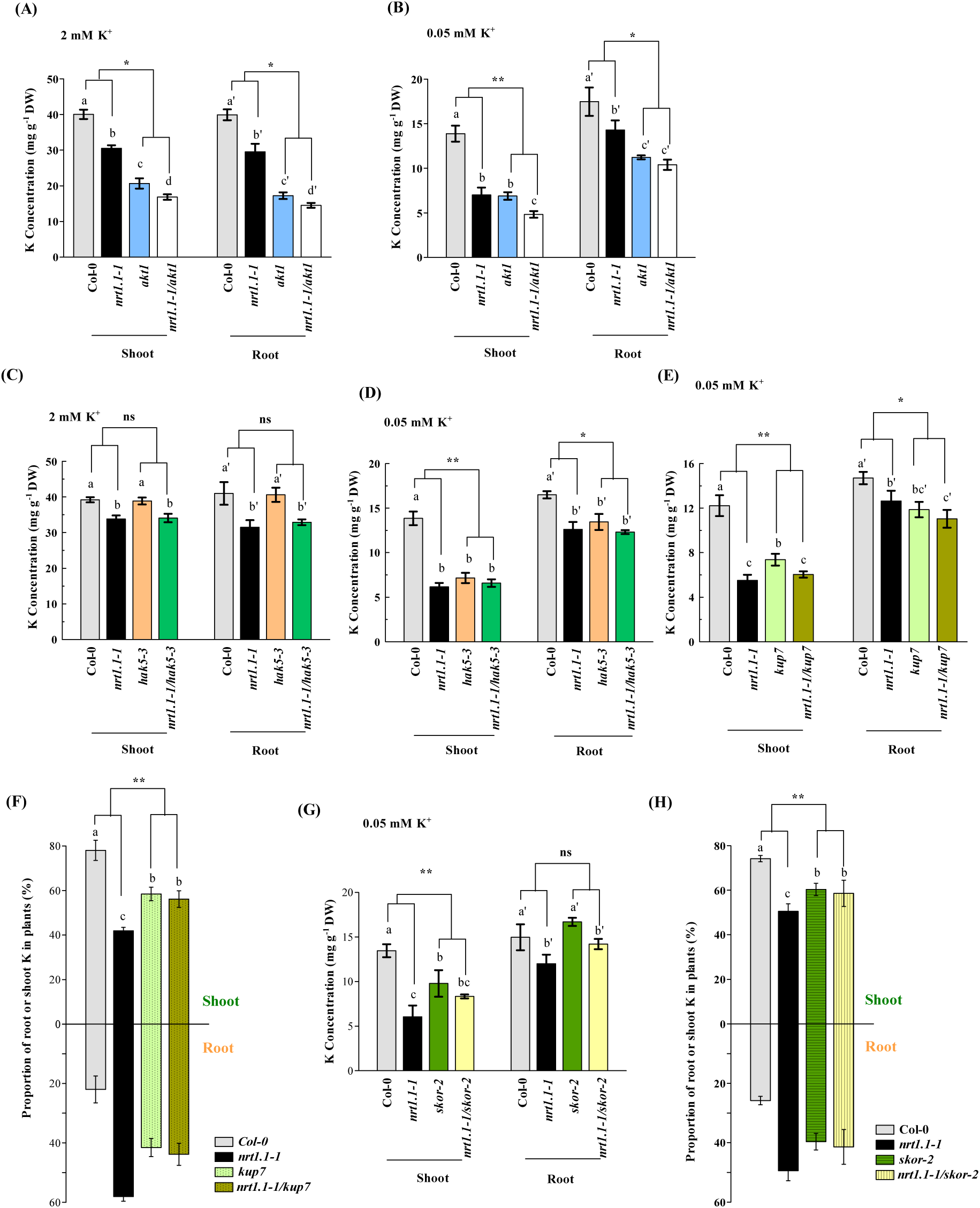
Physiological interactions between NRT1.1 and K^+^ channels. (A) and (B) The K concentrations in Col-0, *nrt1.1-1, akt1*, and *nrt1.1-1*/*akt1* plants. (C) and (D) The K concentrations in Col-0, *nrt1.1-1, hak5-3*, and *nrt1.1-1*/*hak5-3* plants. (E) K concentrations and (F) distributions in Col-0, *nrt1.1-1, kup7*, and *nrt1.1-1*/*kup7* plants. (G) K concentrations and (H) distributions in Col-0, *nrt1.1-1, skor-2*, and *nrt1.1-1*/*skor-2* plants. The 4-d-old seedlings were transferred to an agar medium containing 2 mM K^+^ or 0.05 mM K^+^ for 8 d and subsequently analyzed for K levels, as described in the Materials and Methods section. Bars represent the mean ± SD (n = 5). Different letters above bars indicate significant differences at *P* < 0.05 (LSD test). Asterisks indicate a significant genotype by genotype interaction; ns, non-significant (* *P* < 0.05, ** *P* < 0.01, two-way ANOVA).

## Discussion

K^+^ and NO_3_^-^ are the major forms of potassium and nitrogen that are absorbed by roots of most terrestrial plants. In addition, they, respectively, represent the most abundant inorganic cation and anion sources in plant cells, and are thus also the major forms of potassium and nitrogen that are transported within plants (Du *et al*., 2019). In this study, we revealed a close relationship between NO_3_^-^ and K^+^ homeostasis, which was mediated by the NO_3_^-^ transporter NRT1.1, in the root tissues of *Arabidopsis*.

NRT1.1 was initially characterized as an NO_3_^-^ transporter involved in root NO_3_^-^ uptake from the growth medium (Tsay *et al*., 1993). Further, it was found to be a bidirectional transporter (Remans *et al*., 2006), thereby acting as a component involved in the xylem NO_3_^-^ loading in the root stele, which confers the root-to-shoot allocation of NO_3_^-^ (Léran *et al*., 2013). Several studies have revealed many additional functions such as regulation of another NO_3_^-^ transporter NRT2.1 (Ho *et al*., 2009), auxin transport (Krouk *et al*., 2010), and regulation of tolerance to NH_4_^+^ toxicity (Jian *et al*., 2018). These functions of NRT1.1 are independent of its NO_3_^-^ transport activity, because the absence of NO_3_^-^ supply in the growth medium did not affect the above functions. However, our finding that wild-type plants did not show higher K levels in both shoots and roots than in *nrt1.1* knockout mutants when NH_4_^+^ was the sole nitrogen source shows that the activity of NO_3_^-^ transport across the plasmalemma is probably necessary for NRT1.1 to coordinate K^+^ homeostasis in plants (Fig. S7c,d). Theoretically, NO_3_^-^ transport-coordinated K^+^ homeostasis in plants follows a general mechanism. However, the other nitrate uptake NRTs did not act like NRT1.1 when coordinating K^+^ homeostasis in plants. Previous studies showed that NRT1.1 contributes to > 70% of the total root NO_3_^-^ uptake in NO_3_^-^-sufficient medium (Huang *et al*., 1996; Wang *et al*., 1998), suggesting that the other NRTs (i.e., NRT1.2, NRT2.1, NRT2.2, NRT2.4, and NRT2.5) might just be responsible for < 30% of NO_3_^-^ uptake. Therefore, an insufficient NO_3_^-^ transport of any of these five NRTs might explain why none of them plays an evident role in improving K^+^ homeostasis in plants. This also suggests that the action of NRT1.1 to improve K^+^ homeostasis might require a higher NO_3_^-^ supply to ensure sufficient NO_3_^-^ transport. This notion was supported by the observation that the increase of NO_3_^-^ supply elevated the K levels in wild-type plants that have a normal NRT1.1 function (Fig. 1a,b). In addition, the finding that NRT1.1-null mutants did not have distinguishable growth phenotype and K levels compared with those of wild-type plants in low NO_3_^-^ growth medium (0.2 mM NO_3_^-^; Fig S4) further supports the above notion.

Interestingly, the coordination of K^+^ homeostasis by NRT1.1 in *Arabidopsis*, including root uptake and root-to-shoot allocation processes, depend on the nature of NRT1.1 expression pattern in the root epidermis-cortex and central vasculature, respectively. Hence, determining how such expression pattern of NRT1.1 in root tissues improves the uptake and root-to-shoot allocation of K^+^ becomes necessary. Previously, NRT1.1 and other NRT transporters were shown to be a kind of proton-coupled transporters that transport NO_3_^-^ across the plasmalemma together with an equal amount of proton (H^+^) influx via a symport mechanism (Tsay *et al*., 1993; Fang *et al*., 2016). However, the H^+^ levels in the regular growth media were considerably less than those of NO_3_^-^. Thus, an H^+^ efflux mechanism of root cells should be required to compensate for the consumed H^+^ due to NO_3_^-^ uptake, which consequently ensures the subsequent NO_3_^-^ uptake. The uptake of K^+^ across the plasmalemma by K^+^ channels of the root cortex cells is known to be electrically coupled to H^+^-ATPase-mediated H^+^ efflux (Behl & Raschke, 1987; Wang & Wu, 2013). Since K^+^ is the most abundant inorganic cation absorbed by roots, the amount of H^+^ efflux owing to K^+^ uptake could be a key compensation to the consumed H^+^ by NO_3_^-^ uptake, which might also be a mechanism to retain the charge balance during NO_3_^-^ –K^+^ fluxes across the plasmalemma and thus to ensure a normal electrophysiological environment in the cells. In this context, NRT1.1 is probably coordinated by K^+^ channels in the root epidermis-cortex. Our two-way ANOVA based on K levels in *nrt1.1*/*akt1, nrt1.1*/*hak5*, and *nrt1.1*/*kup7* double mutants and their corresponding monomutants and wild-type plants supported the above notion, that is, the NRT1.1 interplay with K^+^ channels in the root epidermis-cortex could improve the plant K uptake (Fig. 8a,b,c,e). As all the investigated K^+^ channels could physiologically interplay with NRT1.1 to improve K uptake, we suggest that these interplays should function in a nonspecific manner.

The two-way ANOVA based on K distribution in the shoot and root tissues of *nrt1.1/skor* and *nrt1.1*/*kup7* double mutants and their corresponding monomutants and wild-type plants also revealed a physiological interaction between NRT1.1 and K^+^ channels in the root central vasculature, which improves the root-to-shoot allocation of K (Fig. 8e-h). In addition to this mechanism, transcript regulation by NRT1.1 might also play a role in improving the root-to-shoot allocation of K, because the expression of *SKOR* was positively regulated by NRT1.1 (Fig. 7a). We also analyzed the K distribution in the shoot and root tissues of *nrt1.1/akt1* and *nrt1.1/hak5* double mutants and their corresponding monomutants and wild-type plants, but no physiological interaction was noted between NRT1.1 and AKT1 or HAK5 in improving the root-to-shoot allocation of K, as revealed by the two-away ANOVA (Fig. S11c,d), indicating that the K^+^ uptake channels might not be involved in the NRT1.1-coordinated root-to-shoot allocation of K. Notably, the contribution of NRT1.1 to the root-to-shoot allocation of K^+^ is only remarkable under the K-limited condition, but not under the K-sufficient condition (Fig. 5d). Our finding that insufficient NO_3_^-^ supply did not efficiently support the role of NRT1.1 in the uptake and root-to-shoot allocation of K^+^ (Fig. 4e) indicated that a higher ratio of NO_3_^-^ to K^+^ transport across the cell plasmalemma might be a prerequisite for NRT1.1 to remarkably contribute the root-to-shoot allocation of K^+^ in plants. As the K^+^ in plants could be easily allocated to shoots after its root uptake from the growth medium (De Boer, 1999), the amount of K^+^ loading into the root xylem needs to be very higher under the K-sufficient condition, which results in a lower ratio of NO_3_^-^ to K^+^ transport across the cell plasmalemma in the root central vasculature. This might explain why the contribution of NRT1.1 to the root-to-shoot allocation of K^+^ is not remarkable under the K-sufficient condition.

As both the uptake and root-to-shoot allocation of K^+^ were significantly improved by NRT1.1 under the K-limited condition (Figs 3b, 5d), the *nrt1.1* knockout mutants had shorter roots, less biomass, and severer leaf senescence, compared with the wild-type plants in the low-K growth medium (Fig. 2e,f,a), indicating that NRT1.1 is required for plants to resist K deficiency. Owing to the limited availability of K^+^ in most natural soils, plants often suffer from K^+^-deficiency stress (Wang & Wu, 2013). Therefore, plants have to evolve a complex signaling and physiological regulatory network to adapt to K^+^-deficient environments, which helps them survive under K^+^-deficiency stress (Ashley *et al*., 2006; Tsay *et al*., 2011). We found that the activity of NRT1.1-mediated NO_3_^-^ uptake was significantly up-regulated in response to low-K^+^ stress (Fig. 4f). Considering a role of NRT1.1 in improving both root uptake and root-to-shoot allocation of K^+^ in plants, the up-regulation of NRT1.1 activity by low-K^+^ stress could be recognized as an adaption mechanism for plants to help them survive in K^+^-deficient environments. Therefore, for crop cultivation, designing other more practical methods for improving NRT1.1-mediated NO_3_^-^ uptake and increasing the utilization efficiency of K fertilizers is necessary. However, how low-K^+^ stress up-regulates NRT1.1 activity is not yet known. In our study, we observed that the rate of root NO_3_^-^ uptake in low-K medium was clearly less than that in sufficient-K medium (Fig. 4f). Therefore, an insufficient K supply might result in mild nitrogen starvation in plants, which could favor an up-regulation of NRT1.1 to compensate for the decreased NO_3_^-^ uptake. Nevertheless, further studies are required to experimentally verify theis assumption.

In conclusion, we showed that the NO_3_^-^ transport activity of NRT1.1 plays an important role in the K nutrition in plants: its expression in root epidermis-cortex coordinates with the K^+^ uptake channels such as AKT1, HAK5, and KUP7 to improve the root K^+^ uptake from the growth medium, whereas its expression in the root central vasculature coordinates with the channels that load K^+^ into the xylem, such as SORK and KUP7, to facilitate the root-to-shoot allocation of K^+^ (Fig. 9). In the current agricultural practices in China, excessive N and K fertilizers are usually applied by farmers; this reduces the utilization efficiencies of these fertilizers and results in environmental pollution (Guo *et al*., 2010; Zhang, 2017). Our findings indicate that designing practical methods for enhancing NRT1.1 activity in crops might be possible to improve simultaneously the utilization efficiencies of N and K fertilizers in agricultural production.

**Figure 9.**
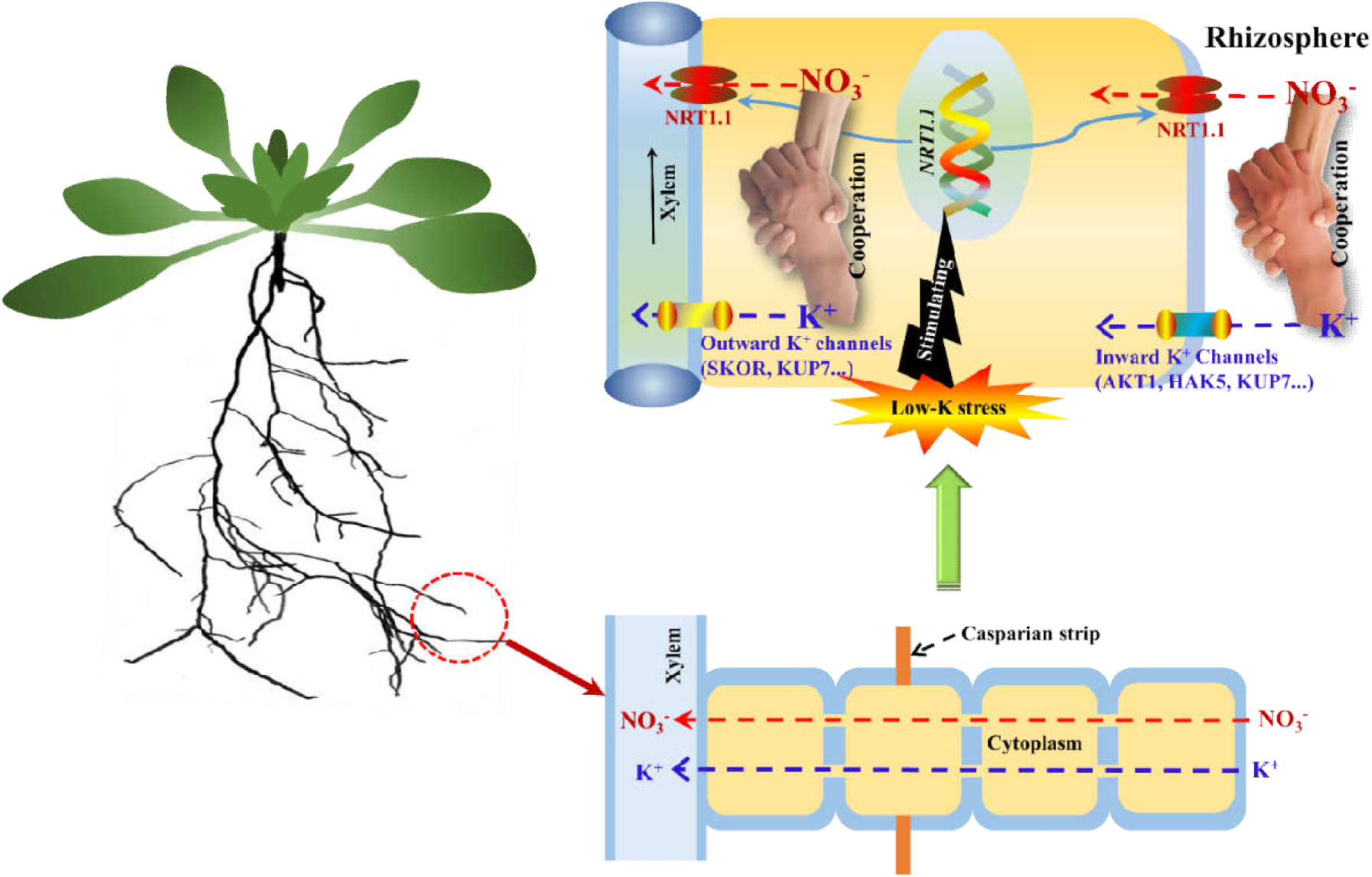
Schematic model of how NRT1.1 responds to low-K stress in *Arabidopsis*. Under insufficient K growth conditions, the expression of the *NRT1.1* gene and its encoding protein in the root epidermis-cortex and central vasculature was induced, which coordinated with the K^+^ uptake channels (such as AKT1, HAK5, and KUP7) and xylem K^+^ loaders (such as SORK and KUP7) to improve root K^+^ uptake from the external environment and facilitate K^+^ root-to-shoot allocation, respectively.

## Acknowledgments

The authors thank Dr. Philippe Nacry for sharing seeds and Dr. Yi Wang (China Agricultural University) for providing the yeast strains. This work was financially supported by the Natural Key R&D Program of China [2016YFD0200103], the Natural Science Foundation of China [31622051 and 31670258], and the Fundamental Research Funds for the Central Universities [2017XZZX002-06].

## Author Contributions

X.Z.F., X.X.L. and C.W. J. conceived the project and designed the experiments; C.C. and X.X.L performed most of the experiments and analyzed data; Y.X.Z. and J.Y.Y. helped with generation of double mutants; X.Z.F. and C.W. J. wrote the article with contributions from all authors.

## Supporting Information

**Fig. S1** Effect of NO_3_^-^ on the amount of K^+^ absorbed per weight of roots in Col-0 plants.

**Fig. S2** Root growth of *Arabidopsis thaliana* Col-0, *nrt1.1-1*, and *chl-5* plants to low-K^+^ stress.

**Fig. S3** Comparisons of growth and K concentration between *Ler* and *chl1-6* plants.

**Fig. S4** Comparisons of growth and K concentrations between Col-0 and *nrt1.1* plants in low nitrate medium.

**Fig. S5** Growth responses of Col-0, *nrt1.2, nrt2.1, nrt2.2, nrt2.5, Ler*, and *nrt2.4* plants to low-K^+^ stress.

**Fig. S6** Comparison of low-K sensitivity between *chl1-5* mutants and complementation line.

**Fig. S7** Comparisons of growth and K concentration between Col-0 and *nrt1.1* plants in nitrate-free medium.

**Fig. S8** The K concentrations in Col-0, *nrt1.2, nrt2.1, nrt2.2, nrt2.5, Ler*, and *nrt2.4* plants.

**Fig. S9** Effect of low-K^+^ stress on the expression of *NRTs* in the root of Col-0 plants.

**Fig. S10** Heatmap analysis of changes in gene expression regulated by K in the roots of Col-0 and *nrt1.1-1* plants.

**Fig. S11** Roles of K^+^ channels in NRT1.1-imrpoved K nutrition is associated with the biological function of K^+^ channels.

**Table S1** Primers used in this study.

