## Supplemental files for "K^+^ uptake and root-to-shoot allocation in *Arabidopsis* require coordination of nitrate transporter1/peptide transporter family member NPF6.3/NRT1.1"

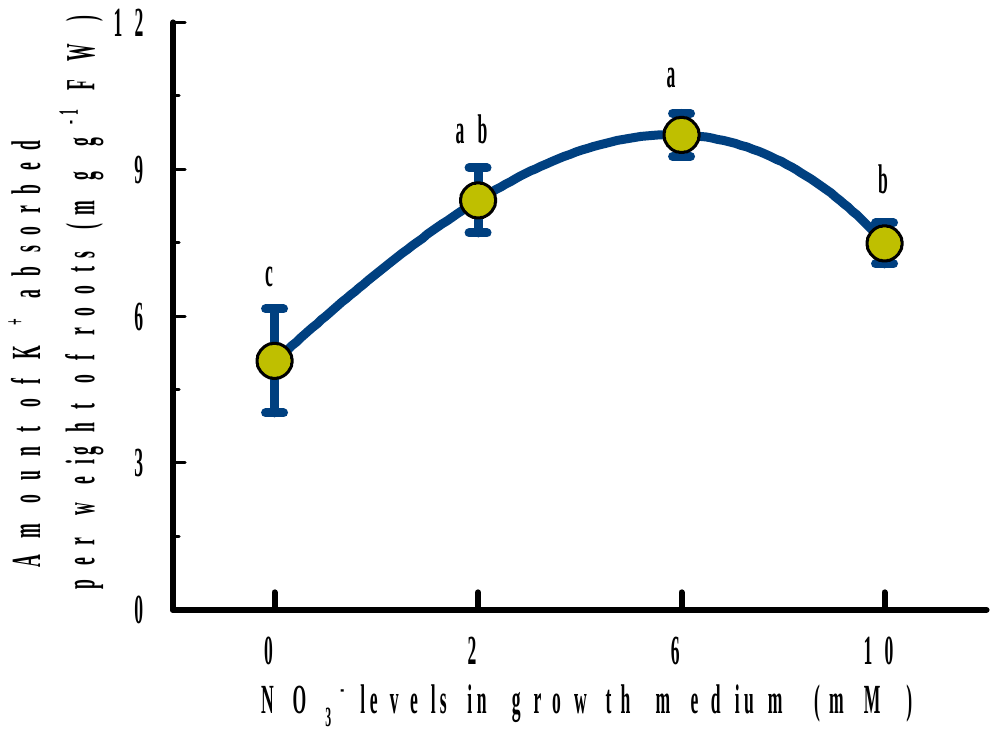

Supplemental Figure 1. Effect of NO_3_^-^ on the amount of K^+^ absorbed per weight of roots in *Arabidopsis thaliana* Col-0 plants. Treatments are the same as those in Figure 1. Bars represent the SD (n = 5). Different letters above bars indicate significant differences at *P* < 0.05 (LSD test).

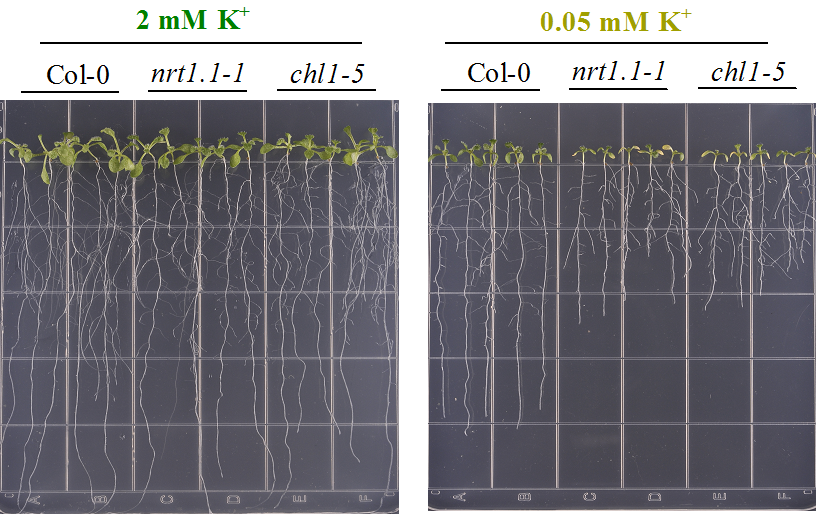

Supplemental Figure 2. Root growth of *Arabidopsis thaliana* Col-0, *nrt1.1-1*, and *chl-5* plants to low-K^+^ stress. The 4-d-old seedlings were treated in the same way as that mentioned in Fig. 2.

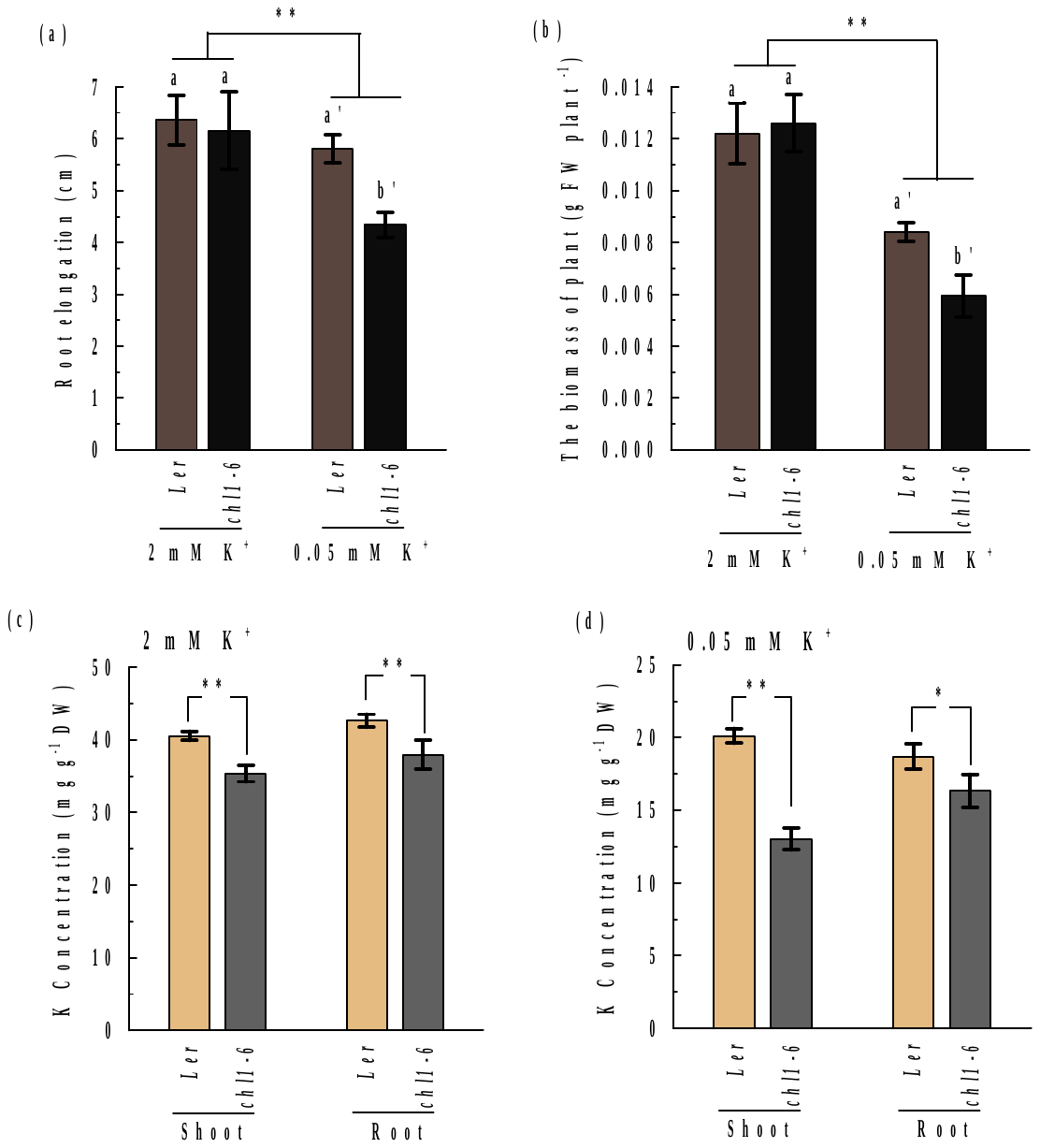

Supplemental Figure 3. Comparisons of growth and K concentration between *Arabidopsis thaliana Ler* and *chl1-6* plants. The 4-d-old seedlings were treated in the same way as that shown in Fig. 2, and then the root elongation (a), biomass (b), and K concentrations of the shoot and root (c, d) of plants were analyzed. Bars represent the mean ± SD (n = 5). (a) and (b) Different letters above bars indicate significant differences at *P* < 0.05 (two-tailed Student’s t-test); asterisks indicate a significant genotype by treatment interaction (** *P* < 0.01, two-way ANOVA). (c) and (d) Asterisks indicate significant differences between genotypes (* *P* < 0.05, ** *P* < 0.01, two-tailed Student’s *t*-test).

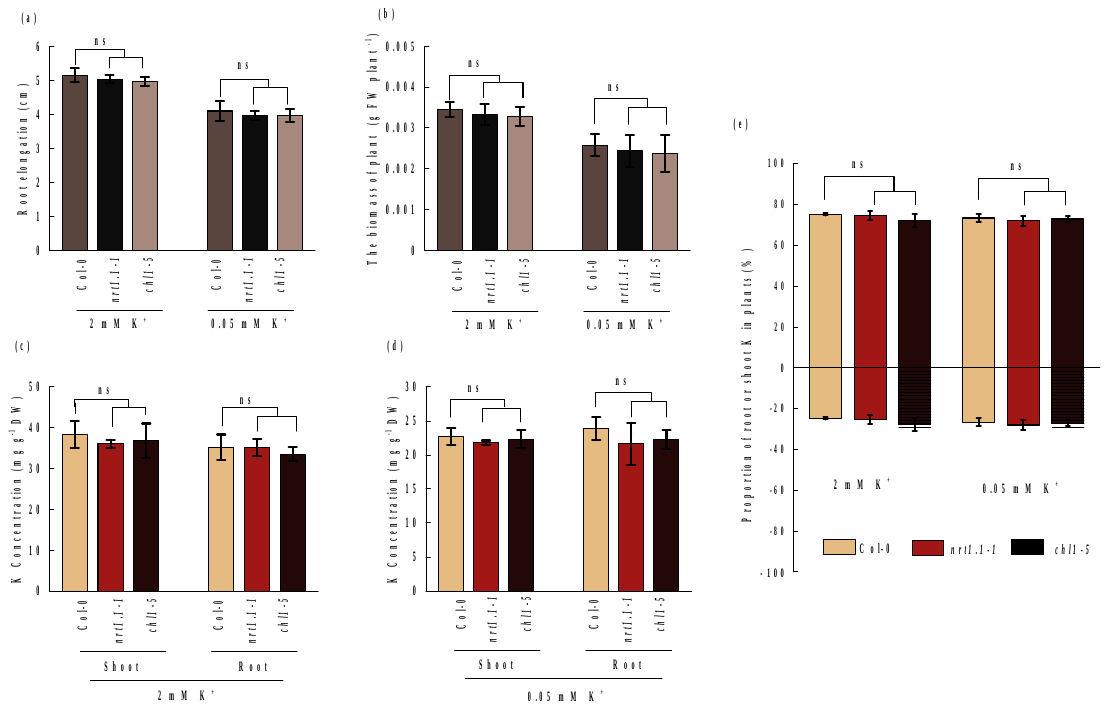

Supplemental Figure 4. Comparisons of growth and K concentrations between *Arabidopsis thaliana* Col-0 and *nrt1.1* plants in low nitrate medium. The 4-d-old seedlings were transferred to a low nitrate nutrient agar (0.2 mM NO_3_^-^) media with 2 mM or 0.05 mM K^+^. The analyses were performed after 8 d of seedling transfer. (a) Root elongation; (b) biomass of plants; (c, d) shoot and root K concentrations of plants under 2 mM and 0.05 mM K^+^ treatment; (e) proportions of K distributed in the shoots and roots. Bars represent the mean ± SD (n = 5); asterisks indicate significant differences compared with the control; ns, non-significant (** *P* < 0.01, two-tailed Student’s *t*-test).

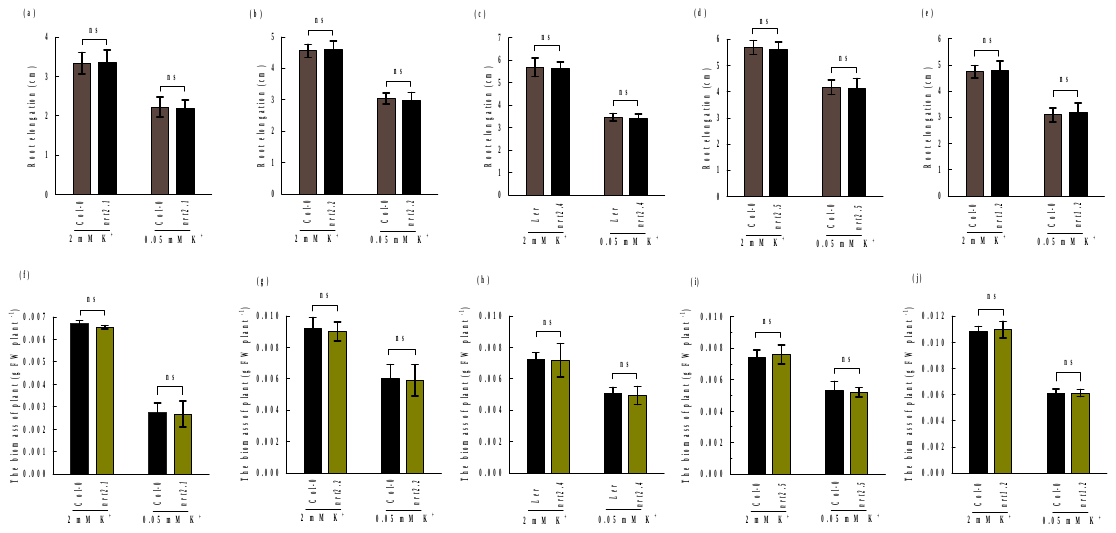

Supplemental Figure 5. Growth responses of *Arabidopsis thaliana* Col-0, *nrt1.2*, *nrt2.1*, *nrt2.2*, *nrt2.5*, *Ler*, and *nrt2.4* plants to low-K^+^ stress. (a−e) Root elongation; (f−j) plant biomass. The 4-d-old seedlings were transferred to a low-nitrate (0.2 mM NO_3_^-^) nutrient media (a−d and f−i) or sufficient-nitrate (6 mM NO_3_^-^) nutrient media (e and j) with 2 mM or 0.05 mM K^+^. The analyses were performed after 8 d of seedling transfer. Bars represent the mean ± SD (n = 4−5); asterisks indicate significant differences between genotypes; ns, non-significant (** *P* < 0.01, two-tailed Student’s *t*-test).

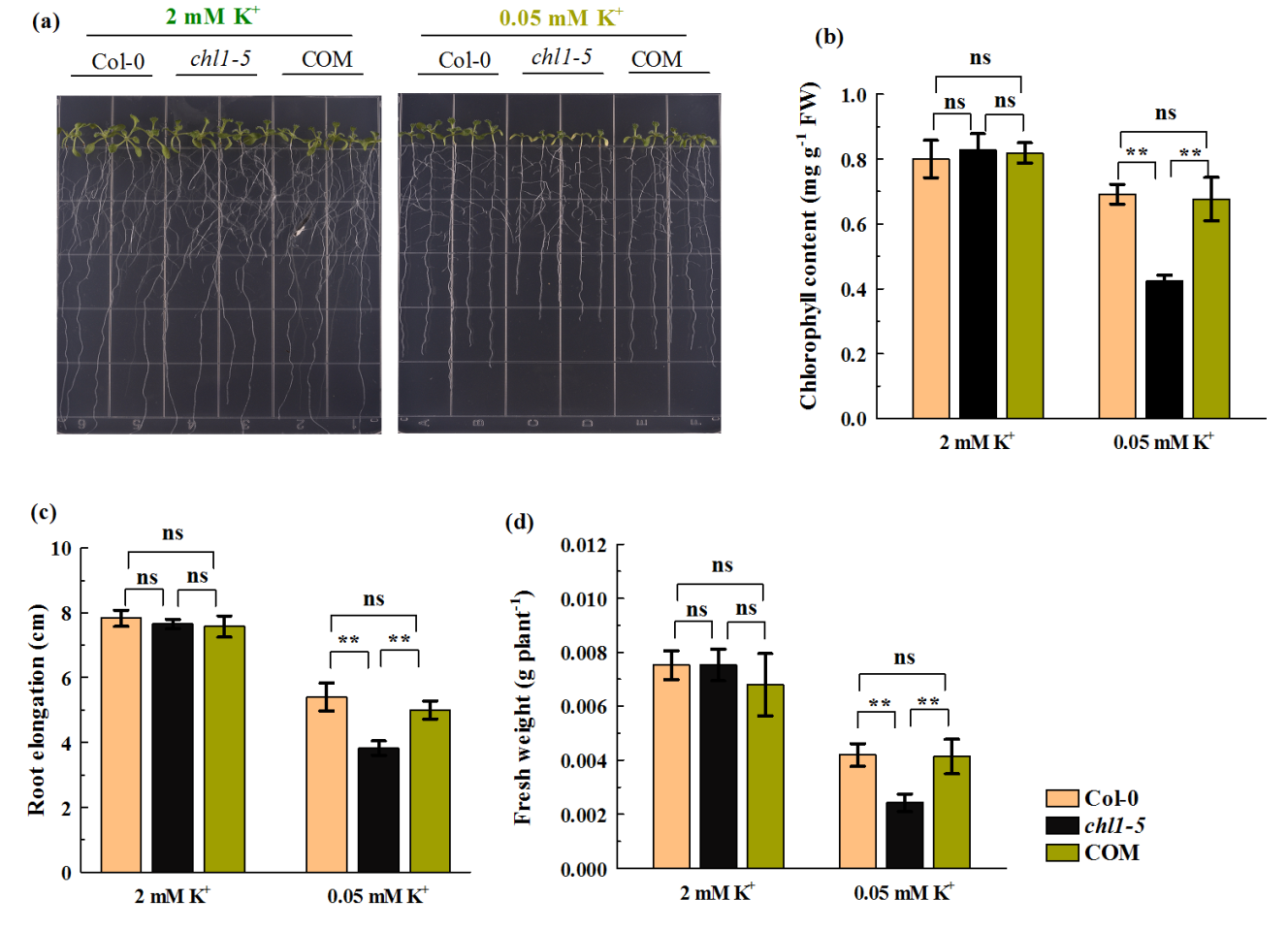

Supplemental Figure 6. Comparison of low-K sensitivity between *chl1-5* mutants and complementation line (COM, *pNRT1.1::NRT1.1-GFP*/*chl1-5*). The 4-d-old seedlings were treated in the same way as that shown in Fig. 2. (a) Root phenotypes; (b) chlorophyll content of leaves; (c) root elongation; (d) biomass of plants. Bars represent the mean ± SD (n = 4−5); asterisks indicate significant differences between genotypes; ns, non-significant (** *P* < 0.01, two-tailed Student’s *t*-test).

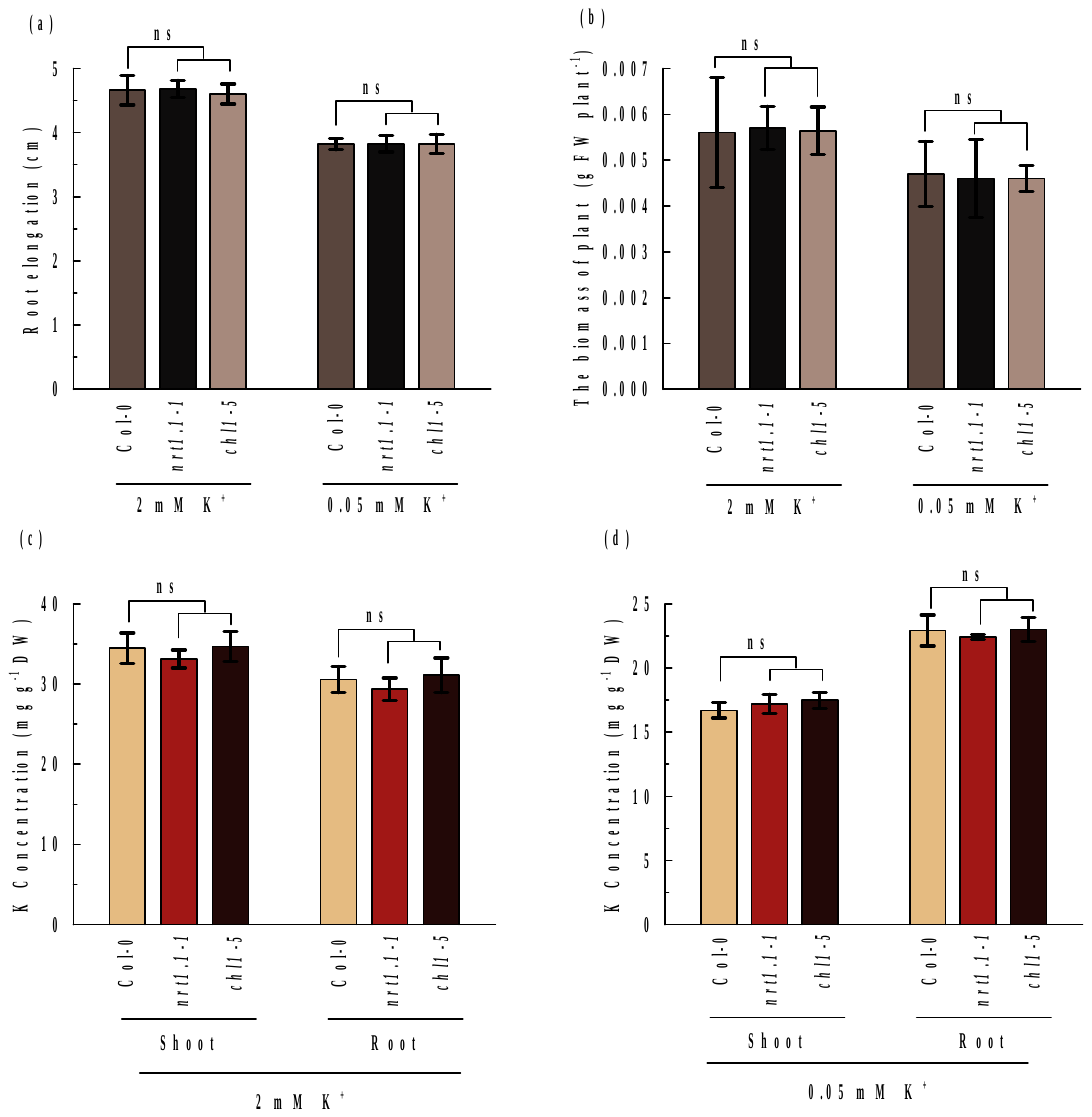

Supplemental Figure 7. Comparisons of growth and K concentration between *Arabidopsis thaliana* Col-0 and *nrt1.1* plants in nitrate-free medium. The 4-d-old seedlings were transferred to a nitrate-free nutrient agar media with 2 mM or 0.05 mM K^+^. The analyses were performed after 8 d of seedling transfer. (a) Root elongation; (b) biomass of plants; (c) and (d) K concentrations in plants under 2 mM and 0.05 mM K^+^ treatments, respectively. Bars represent the mean ± SD (n = 5); asterisks indicate significant differences between genotypes; ns, non-significant (** *P* < 0.01, two-tailed Student’s *t*-test).

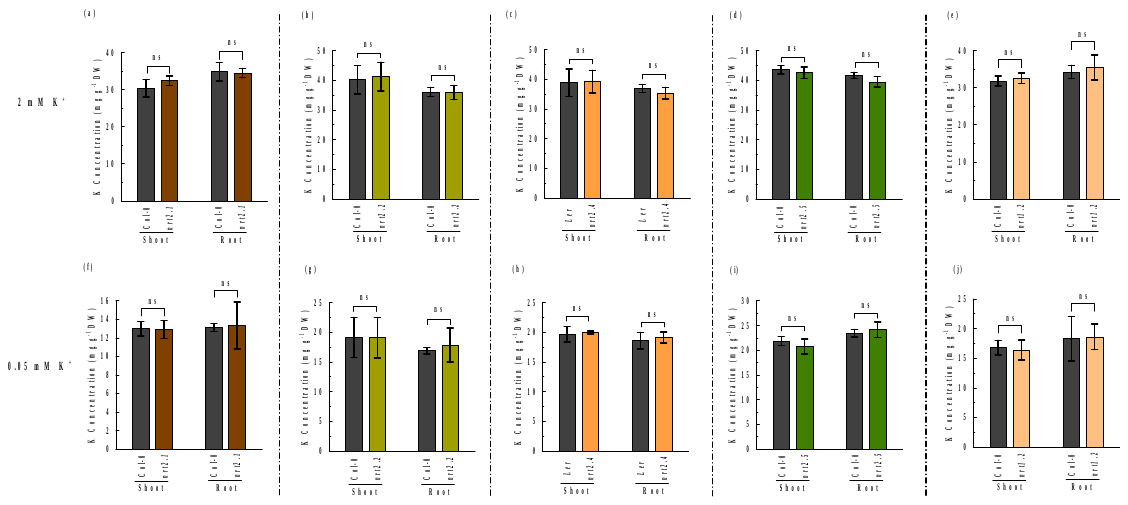

Supplemental Figure 8. The K concentrations in *Arabidopsis thaliana* Col-0, *nrt1.2*, *nrt2.1*, *nrt2.2*, *nrt2.5*, *Ler*, and *nrt2.4* plants. (a−e) The K concentrations of plants under 2 mM K^+^ treatment; (f−j) The K concentrations of plants under 0.05 mM K^+^ treatment. The 4-d-old seedlings were treated in the same way as that mentioned in Fig. S4 (a−d and f−i) or Fig. 2 (e and j). The 4-d-old seedlings were transferred to a low-nitrate (0.2 mM NO_3_^-^) nutrient media (a−d and f−i) or sufficient-nitrate (6 mM NO_3_^-^) nutrient media (e and j) with 2 mM or 0.05 mM K^+^. The analyses were performed after 8 d of seedling transfer. Bars represent the mean ± SD (n = 5); ns, non-significant (** *P* < 0.01, two-tailed Student’s *t*-test).

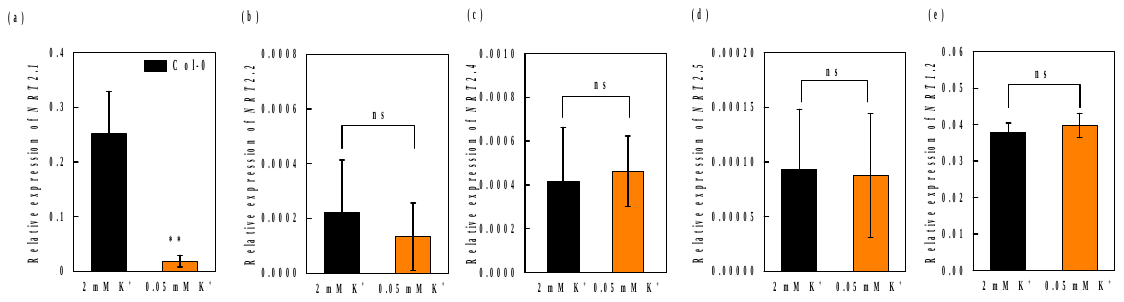

Supplemental Figure 9. Effect of low-K^+^ stress on the expression of *NRTs* in the root of *Arabidopsis thaliana* Col-0 plants. The 4-d-old seedlings were transferred to agar media containing 2 mM K^+^ or 0.05 mM K^+^. The analyses were performed after 5 d of seedling transfer. Relative expressions of *NRT2.1* (a), *NRT2.2* (b), *NRT2.4* (c), *NRT2.5* (d), and *NRT1.2* (e). Relative expression levels were normalized to those of *UBQ10*. Bars represent the mean ± SD (n = 12); asterisks indicate significant differences between treatments; ns, non-significant (** *P* < 0.01, two-tailed Student’s *t*-test).

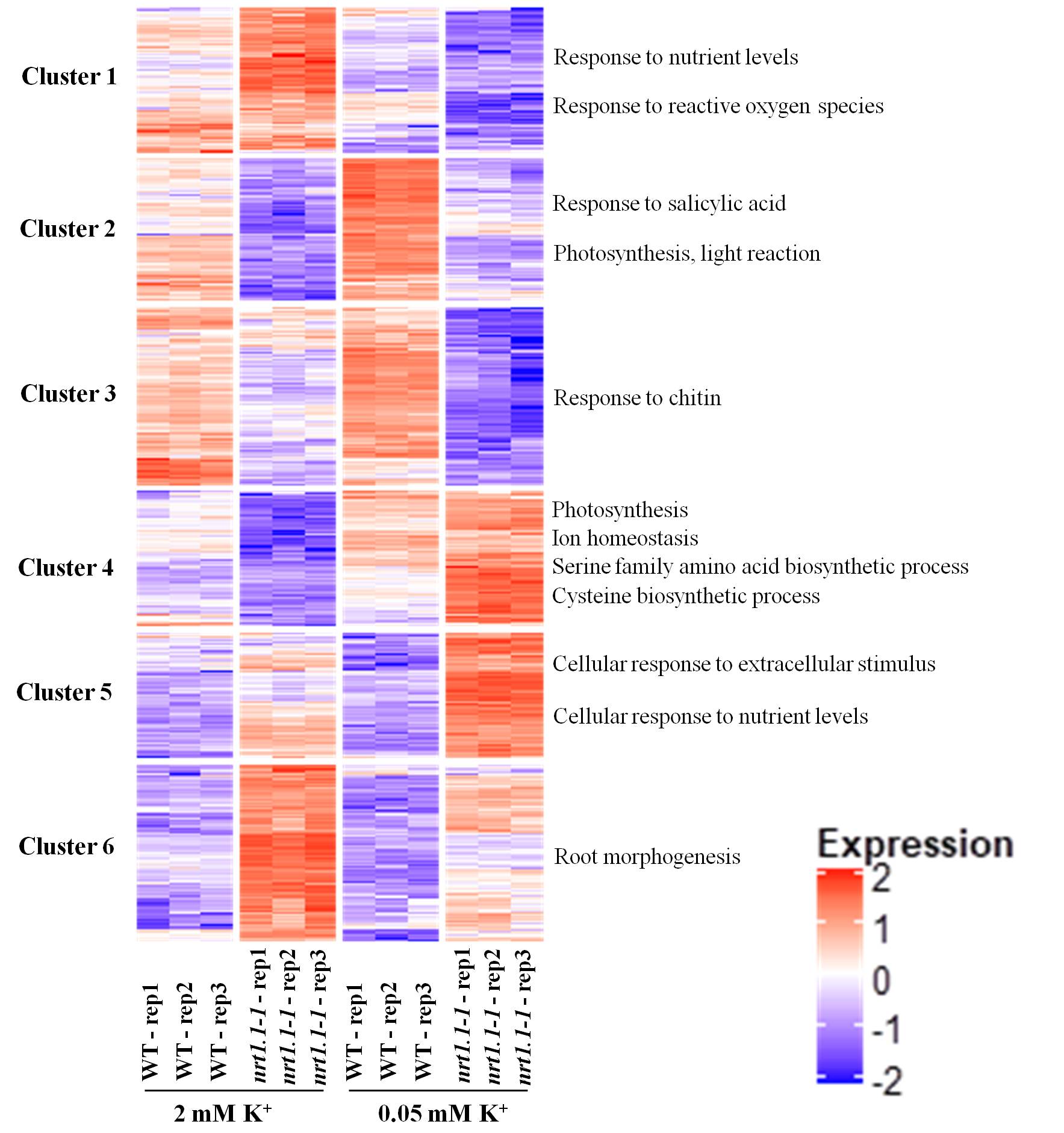

Supplemental Figure 10. Heatmap analysis of changes in gene expression regulated by K in the roots of Arabidopsis Col-0 and *nrt1.1-1* plants. The 4-d-old seedlings were transferred to a media with 2 mM or 0.05 mM K^+^ for 5 d, and roots were collected to perform transcriptome analysis, as described in the Materials and Methods section. Changes in response to K treatment with a *p*-value of <0.05 were selected for clustering analysis.

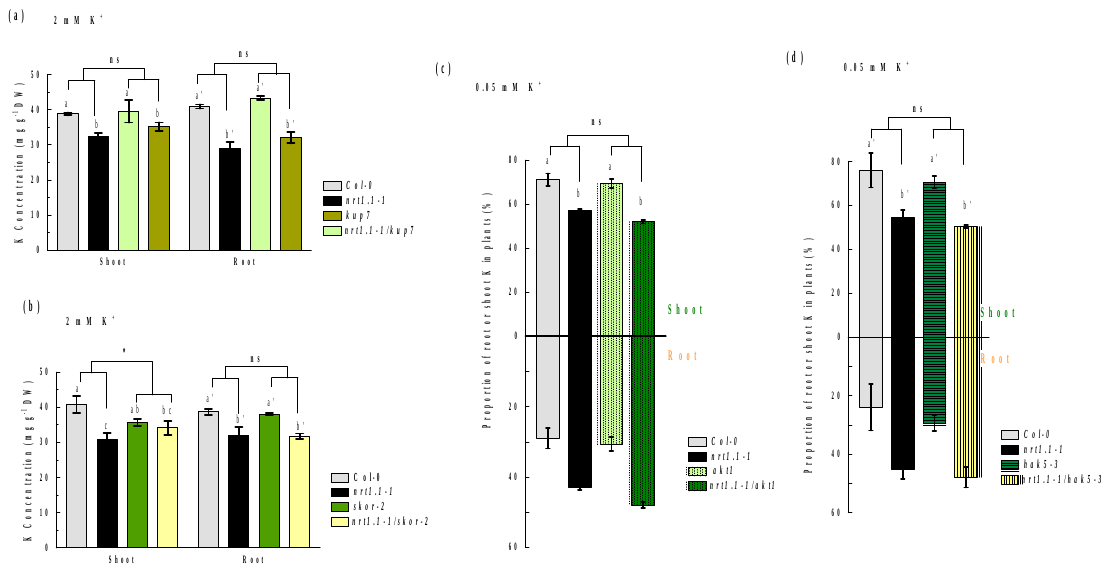

Supplemental Figure 11. Roles of K^+^ channels in NRT1.1-imrpoved K nutrition is associated with the biological function of K^+^ channels. (a) K concentrations in the shoots and roots of Col-0, *nrt1.1-1*, *kup7*, and *nrt1.1-1*/*kup7* plants; (b) K concentrations in the shoots and roots of Col-0, *nrt1.1-1*, *skor-2*, and *nrt1.1-1*/ *skor-2* plants; (c) K distributions in Col-0, *nrt1.1-1*, *akt1*, and *nrt1.1-1*/*akt1* plants; (d) K distributions in Col-0, *nrt1.1-1*, *hak5-3*, and *nrt1.1-1*/*hak5-3* plants. The 4-d-old seedlings were transferred to an agar medium containing 2 mM K^+^ or 0.05 mM K^+^ for 8 d and subsequently analyzed for K levels, as described in the Materials and Methods section. Bars represent the mean ± SD (n = 5); different letters above bars indicate significant differences at *P* < 0.05 (LSD test); asterisks indicate a significant genotype by genotype interaction; ns, not significant (* *P* < 0.05, two-way ANOVA).

**Supplemental Table 1.** Primers used in this study.

| Gene | Primer | Sequence (5ʹ-3ʹ) | Note |
| --- | --- | --- | --- |
| Real-time qPCR | | | |
| AT4G05320 | *UBQ10* F | ACCCTAACGGGAAAGACGA | qPCR |
|  | *UBQ10* R | GGAGCCTGAGAACAAGATGAA |  |
| AT1G12110 | *NRT1.1* F | TTCTCGTGACAATCGTCG | qPCR |
|  | *NRT1.1* R | TGGAATACTCGGCTCATCAT |  |
| AT1G69850 | *NRT1.2* F | GGATTCGGTGTTTCTACCAT | qPCR |
|  | *NRT1.2* R | CGCAATGATTTGAGGGAC |  |
| AT1G08090 | *NRT2.1* F | AACAAGGGCTAACGTGGATG | qPCR |
|  | *NRT2.1* R | CTGCTTCTCCTGCTCATTCC |  |
| AT1G08100 | *NRT2.2 F* | CAGGTGGAAACAGAGCTGCCATGG | qPCR |
|  | *NRT2.2 R* | GACCATAGATACAACGGCAGTGACGAG |  |
| AT5G60770 | *NRT2.4* F | CCGTCTTCT CCA TGTCTTTC | qPCR |
|  | *NRT2.4* R | CTGACCATTGAACATTGTG |  |
| AT1G12940 | *NRT2.5* F | CTCCTCCCTGTTATCCGTGAAA | qPCR |
|  | *NRT2.5* R | AGACGAAAGTGGCGAGAGAGAA |  |
| Mutant screening | | | |
| AT1G12110 | salk_097431 LP | ATATTGGAATCCCTTTCTCGG | Genotyping |
|  | salk_097431 RP | GAGGAAGCGTTTTGACTGTTG |  |
| [AT5G60770](https://www.arabidopsis.org/servlets/TairObject?type=locus&id=132626) | cs27332 LP | AAACTTCTTTGCCCGTCC | Genotyping |
|  | cs27332 RP | ATACCCTTTCGCTTCTCGG |  |
| [AT1G69850](https://www.arabidopsis.org/servlets/TairObject?type=locus&id=136536) | [cs859605](https://www.arabidopsis.org/servlets/TairObject?id=1009287901&type=germplasm) LP | CCACGTCAAGAAGAAGCTTTG | Genotyping |
|  | [cs859605](https://www.arabidopsis.org/servlets/TairObject?id=1009287901&type=germplasm) RP | AAAATATTTGGGCCTCGTGAC |  |
| AT1G08100 | salk _043543 LP | CTAGCGTGAGCACCAAGATTC | Genotyping |
|  | salk _043543 RP | AATGAGTTCACGATGTGGTGC |  |
| AT1G12940 | GK-213H10.06 LP | TCTTGTCACAGTTTGCCCG | Genotyping |
|  | GK-213H10.06 RP | TTCTGAGCCTGATGGTTCG |  |
| AT1G08090 | cs859604 LP | GCAAGCGACTATCATCACTCC | Genotyping |
|  | cs859604 RP | GTTCTCCATGAGCTTCGTGAG |  |
| AT2G26650 | salk _071803 LP | TCCATGTCAAGCTAAGAAGACG | Genotyping |
|  | salk _071803 RP | TCGGTGATAGATAGTGGTGGC |  |
| AT4G13420 | salk _130604 LP | CATAGCTTTTTGTCACTTTGATTC | Genotyping |
|  | salk _130604 RP | TTGTTGAGTTTACTTTGGCCG |  |
| AT3G02850 | cs2103489 LP | TATGAACCGAAACAAACTCGG | Genotyping |
|  | cs2103489 RP | ACACGATCATTCCCATTCTTG |  |
| AT5G09400 | cs805085 LP | ATATCCACTGCTCTCAGACCG | Genotyping |
|  | cs805085 LP | AACAGAGCCTGAGAGGTTATCG |  |
| pBIN-pROK2 T-DNA | Lab1.3 | ATTTTGCCGATTTCGGAAC | Genotyping |
|  | GABI | CCATTTGGACGTGAATGTAGACAC |  |
| Yeast complementation | | |  |
| NRT1.1 | NRT1.1 F | tcgactagtggatcccccgggATGTCTCTTCCTGAAACTAAATCTGATG | (XmaI) |
| NRT1.1 | NRT1.1 R | gataagcttgatatcgaattcTCAATGACCCATTGGAATACTCG | (EcoRI) |
| Transgenic construction | | | |
| Promoter of PHO1 | PHO1 F | tatgaccatgattacgaattcGGTTTACTTATTTTTTCATAGAAATACTTACAT | (EcoRI) |
| Promoter of PHO1 | PHO1 R | ggaagagacatCGTCGCATATAATTTCTTCCGTT |  |
| NRT1.1 | NRT1.1 F | tatgcgacgATGTCTCTTCCTGAAACTAAATCTGATG |  |
| NRT1.1 | NRT1.1 R | gcccttgctcaccatgtcgac ATGACCCATTGGAATACTCGGC | (SalI ) |
| Promoter of Sultr1;2 | Sultr1;2 F | tatgaccatgattacgaattcCATCGGATATATATTTAAGTGTAGTTGGTC | (EcoRI) |
| Promoter of Sultr1;2 | Sultr1;2 R | caggaagagacatAGCTATGTAACTCTGCAAACAGAACA |  |
| NRT1.1 | NRT1.1 F | catagctATGTCTCTTCCTGAAACTAAATCTGATG |  |
| NRT1.1 | NRT1.1 R | GcccttgctcaccatgtcgacATGACCCATTGGAATACTCGGC | (SalI ) |
